# Comprehensive analysis of platelet glycoprotein Ibα glycosylation

**DOI:** 10.1101/2022.07.19.500646

**Authors:** Marie A Hollenhorst, Katherine H Tiemeyer, Keira E Mahoney, Kazuhiro Aoki, Mayumi Ishihara, Sarah C. Lowery, Valentina Rangel-Angarita, Carolyn R Bertozzi, Stacy A Malaker

## Abstract

**Background:** Platelet glycoprotein (GP) Ibα is the major ligand-binding subunit of the GPIb-IX-V complex that binds von Willebrand Factor (VWF). GPIbα is heavily glycosylated, and its glycans have been proposed to play key roles in platelet clearance, VWF binding, and as target antigens in immune thrombocytopenia syndromes. Despite its importance in platelet biology, the glycosylation profile of GPIbα is not well characterized.

**Objectives:** The aim of this study was to comprehensively analyze GPIbα amino acid sites of glycosylation (glycosites) and glycan structures.

**Methods:** GPIbα ectodomain that was recombinantly expressed or that was purified from human platelets was analyzed by Western blot, mass spectrometry (MS) glycomics, and MS glycoproteomics to define glycosites and the structures of the attached glycans.

**Results:** We identified a diverse repertoire of N- and O-glycans, including sialoglycans, Tn antigen, T antigen, and ABH blood group antigens. In the analysis of the recombinant protein, we identified 62 unique O-glycosites. In the analysis of the endogenous protein purified from platelets, we identified at least 48 unique O-glycosites and 1 N-glycosite. The GPIbα mucin domain is densely O-glycosylated. Glycosites are also located within the macroglycopeptide domain and mechanosensory domain (MSD).

**Conclusions:** This comprehensive analysis of GPIbα glycosylation lays the foundation for further studies to determine the functional and structural roles of GPIbα glycans.

**Essentials:**

- Glycosylation of glycoprotein Ibα (GPIbα) is important for platelet function.
- We report a comprehensive and site-specific analysis of human GPIbα glycosylation.
- GPIbα carries sialoglycans, Tn antigen, T antigen, and ABO blood group (ABH) antigens.
- We experimentally determined 48 O-glycosites and 1 N-glycosite by mass spectrometry.

## 1. Introduction

Adhesion of platelets to the subendothelium is a critical event in the hemostatic response to vascular injury. Platelet adhesion is mediated by an interaction between the platelet membrane glycoprotein Ibα (GPIbα) and Von Willebrand Factor (VWF) that is immobilized on the vessel wall.(*1-3*) The importance of this interaction is underscored by pathogenic GPIbα mutations that lead to bleeding in patients with Bernard-Soulier syndrome and platelet-type Von Willebrand disease (VWD).(*4, 5*) In addition to its role in binding VWF, GPIbα regulates platelet clearance and can be a target of pathogenic antibodies in immune-mediated thrombocytopenia syndromes.(*6-8*)

GPIbα is heavily glycosylated; glycans account for approximately 40-60% of the mass of the GPIbα ectodomain.(*9-12*) GPIbα is known to carry both of the two major types of protein glycans: 1) extracellular mucin-type O-glycans and 2) N-glycans.(*9-12*) Mucin-type O-glycosylation is initiated by the addition of *N*-acetylgalactosamine (GalNAc) in α-linkage to serine (Ser) or threonine (Thr) to form the Tn antigen (GalNAcαSer/Thr) (Figure 1A-B).(*13*) Addition of galactose (Gal) in β1-3 linkage to GalNAc forms the unsubstituted core 1 O-glycan (Galβ1-3GalNAcαSer/Thr), also referred to as the T antigen. The unsubstituted core 2 O-glycan [GlcNAcβ1-6(Galβ1-3)GalNAcαSer/Thr] is formed by addition of *N*-acetylglucosamine (GlcNAc) to the core 1 O-glycan. Unsubstituted core 1 and core 2 O-glycans can be extended to form a variety of complex O-glycan structures. N-glycans modify asparagine (Asn) residues and can be one of three major types: high mannose (Man), hybrid, or complex (Figure 1C). O- and N-glycans can terminate with a variety of epitopes including sialic acid (SA) residues and the antigens that define the ABO blood groups (termed the ABH antigens) (Figure 1D).

**Figure 1.**
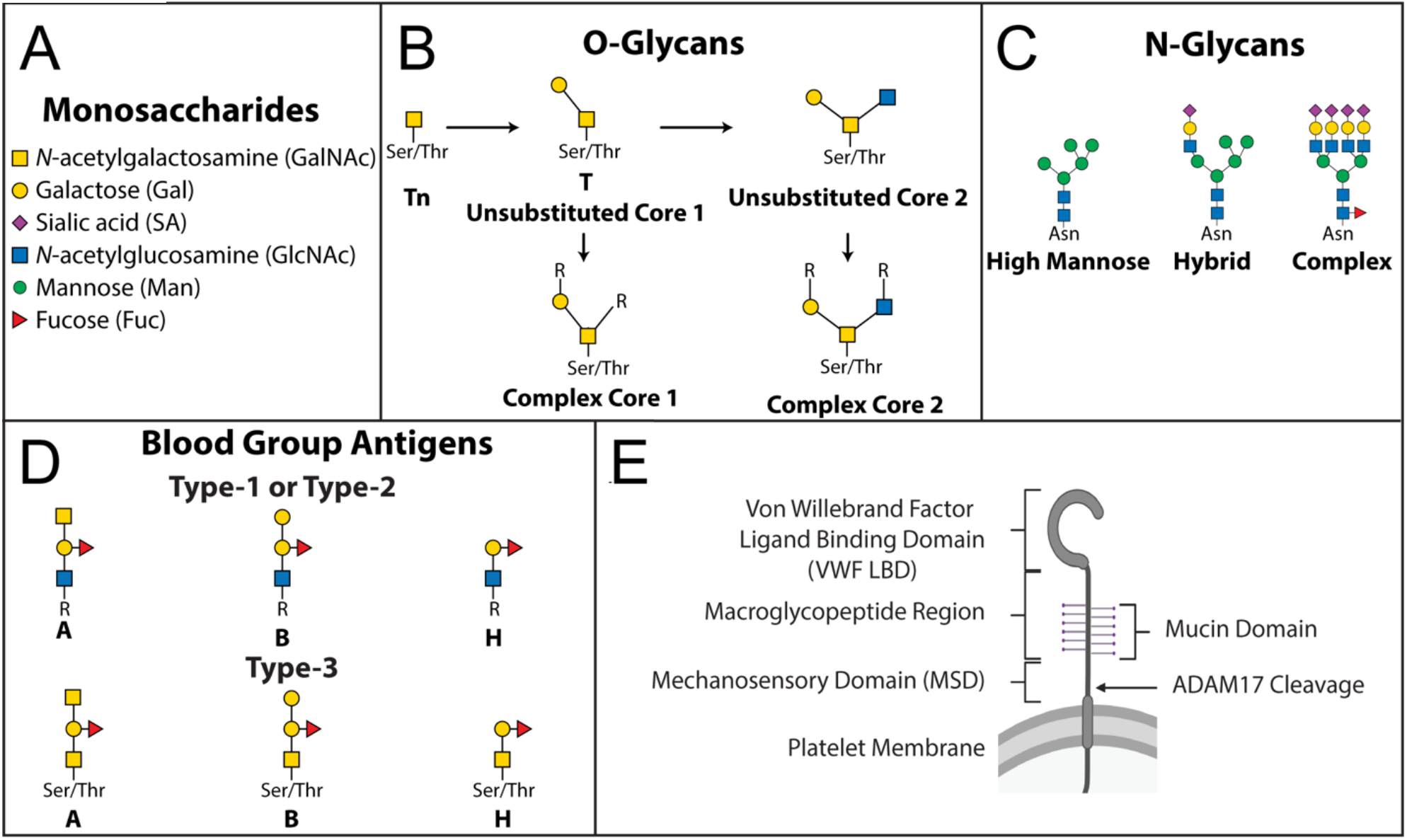
Representative glycan structures. A) Glycans are monosaccharide polymers. Monosaccharides are represented as colored shapes according to Symbol Nomenclature for Glycans (SNFG) standards.(*48*) B) O-glycans covalently modify serine (Ser) or threonine (Thr). Important O-glycans include the Tn and T antigens. Complex core 1 or core 2 O-glycans have variable R groups which can include for example SA. C) N-glycans covalently modify asparagine (Asn). Representative glycans are shown from the three major classes of N-glycans: high mannose, hybrid, and complex. D) ABO blood group is defined by the glycan antigens A, B, and H (H is the antigen found in blood type O individuals). Type-1 and type-2 ABH antigens differ with respect to the linkage between the GlcNAc and Gal (β1,3 for type-1 and β1,4 for type-2). Type-1 ABH antigens are most commonly found as terminal structures on complex O-glycans, and type-2 are most commonly found as terminal structures on N-glycans. Type-3 ABH antigens, also known as mucin-associated ABH antigens, are direct modifications of Ser/Thr.(*13*) E) Domain organization of GPIbα. GPIbα has an N-terminal Von Willebrand Factor (VWF) ligand binding domain (LBD). C-terminal to this is the macroglycopeptide region, which encompasses the mucin domain. Further C-terminal is the mechanosensory domain (MSD). The ADAM17 protease cleaves intact GPIbα to generate the ectodomain; the cleavage site is within the MSD.

GPIbα has been described as a mucin-domain glycoprotein.(*14*) These proteins are characterized by regions rich in O-glycosylated Ser/Thr residues which cause the protein to adopt a rigid “mucin fold” wherein O-glycans extend out like “bottle-brush” bristles from an elongated peptide core.(*15, 16*) The GPIbα mucin domain is embedded within a larger region termed the macroglycopeptide domain (Figure 1E, Figure S1, Table S1).^1^ The O-glycans in the mucin and macroglycopeptide domains are thought to promote projection of the N-terminal VWF ligand binding domain away from the platelet surface.(*15, 16*) GPIbα is endogenously cleaved by the protease ADAM17, resulting in soluble GPIbα ectodomain, also known as glycocalicin.(*17*)

Beyond their structural role in rigidifying the macroglycopeptide region, GPIbα glycans are likely involved in several other aspects of platelet biology. It is well-established that loss of SA from platelet glycans leads to accelerated platelet clearance.(*6, 7, 18-20*) This desialylation has been suggested to cause or contribute to development of thrombocytopenia in several disease contexts, including immune thrombocytopenia (ITP), sepsis, and influenza virus infection.(*7, 20-22*) While the mechanism by which desialylated platelets are cleared is the subject of controversy in the literature, most of the data point towards a central role of GPIbα glycan desialylation.(*6, 7, 18, 19*)

In addition to their likely role in promoting clearance of desialylated platelets, GPIbα glycans are potential antigens targeted by anti-platelet (or anti-megakaryocyte) immune responses in ITP and other immune-mediated thrombocytopenias.(*18-20*) In particular, immune responses against Tn and T antigens have been proposed to be important mechanisms in immune thrombocytopenia syndromes.(*23-25*) Recently, there has been growing interest in the possibility that platelet ABH antigens are relevant to hemostasis, and this has been proposed as a mechanism that could explain the association between ABO blood type and risk of thrombosis or bleeding.(*26, 27*) A final aspect of platelet biology where glycans are likely important is in the unfolding of the GPIbα mechanosensory domain (MSD).(*28*) The MSD unfolds upon VWF binding and pulling, leading to conformational changes in the GPIb-IX complex that ultimately trigger intra-platelet signaling.(*28*) Sialylation of the MSD has also been shown to regulate MSD unfolding.(*28*)

To date, the structural characterization of GPIbα glycans has been limited. GPIbα was proposed to have approximately 60 O-glycosites based on compositional analysis of GPIbα monosaccharides and amino acids.(*10*) However, only 10 glycosites have been previously experimentally determined (Table S2).(*3, 11, 29-31*) Studies of purified GPIbα glycans carried out in the 1980s have revealed the structures of approximately 12 glycans, including core 1 and core 2 O-glycans and complex N-glycans (Figure S2).(*11, 12, 32-35*) Both N- and O-glycans with terminal α(1,2)-linked fucose have been observed.(*33*) These structures are consistent with H antigens, but were not described as such in these publications.(31, 33) Antigen capture enzyme-linked immunosorbent assay (ELISA) and Western blot experiments have suggested that GPIbα bears A and B antigens, but the corresponding glycan structures and glycosites have not been confirmed experimentally.(*36-38*) This knowledge gap hinders our ability to understand the roles of GPIbα glycans in health and disease.

It is not surprising that our knowledge of GPIbα glycosylation is incomplete, as mucin-domain glycoproteins are notoriously challenging to study.(*39*) This is partially due to their resistance to cleavage by standard proteases that are used to generate glycopeptides for MS analysis.(*39*) To address this challenge, we previously characterized secreted protease of C1 esterase inhibitor (StcE), an enzyme that cleaves mucins (i.e., a mucinase) into glycopeptides amenable to MS analysis.(*39*) We demonstrated that StcE proteolyzed GPIbα, suggesting that a mucinase digestion strategy may be effective in facilitating its analysis by MS glycoproteomics.(*39*)

In this study, we undertook a detailed analysis of human GPIbα glycosylation. We purified the GPIbα ectodomain from human platelets and then used a combination of experimental strategies, including mucinase digestion methods, to carry out a glycomics-informed glycoproteomics analysis.(*40*)

## 2. Materials and Methods

### 2.1 Materials

Recombinant GPIbα was obtained from R & D Systems; the gene coding for the human GPIbα ectodomain was overexpressed in a NS0-derived mouse myeloma cell line. Apheresis platelets of blood type A, B, or O were purchased as research products from Bloodworks Northwest (Table S3). Platelets were stored at room temperature with gentle agitation and were used up to 7 days past their transfusion outdate. Further details are available in the Supporting Information.

### 2.2 Purification of GPIbα ectodomain from human platelets

The soluble ectodomain of GPIbα (glycocalicin) was purified from apheresis platelets using an adaptation of previously reported protocols.(*41, 42*) Three to five units of apheresis platelets, generally from distinct donors, were pooled for each purification, so GPIbα in each preparation came from up to 5 different individuals. Platelets were centrifuged and washed with ice-cold PBS. The resulting pellets were resuspended in cold buffer. Platelet suspensions were sonicated and then incubated at 37 °C for 1 hour. After incubation, platelet suspensions were centrifuged and the supernatant was purified by wheat germ agglutinin (WGA) agarose gravity column followed by anion exchange chromatography using a HiTrap DEAE-FF 1 mL column (Cytiva) on an AKTA pure liquid chromatography system (Cytiva). Further details are available in the Supporting Information.

### 2.3 Western blot analysis of GPIbα

Purified GPIbα (5 µg) was separated by sodium dodecyl sulfate-polyacrylamide gel electrophoresis (SDS-PAGE) then transferred to a nitrocellulose membrane. The following primary antibodies were used: anti-A or anti-B murine monoclonal blend (1:10, Ortho-Clinical Diagnostics), rabbit monoclonal anti-GPIbα (1:10,000, Abcam ab 210407, EPR19204). Blots were incubated with the following secondary antibodies for 1 hour at room temperature: IRDye 800 CW goat anti-mouse IgM (LiCOR, 1:15,000), IRDye 800 CW goat anti-rabbit (Li-COR, 1:15,000). Blots were imaged on the Odyssey DLx imaging system (Li-COR).

H antigen was visualized using biotinylated *Ulex europaeus* I (UEA-I, Vector Laboratories). A one hour, room temperature incubation was performed for the secondary antibody (IRDye 800 CW Streptavidin, 1:1000 in Carbo-Free blocking buffer containing 0.2% Tween-20. SA-containing GPIbα glycans were also assessed by lectin blotting using the *Sambucus nigra* (SNA/EBL) biotinylated lectin (Vector Laboratories) at a working concentration of 2 µg/µL. Further details are available in the Supporting Information.

### 2.4 Release of glycans from purified platelet-derived human GPIbα ectodomain for glycomics analysis

Glycan release was performed according to previously reported methods.(*40, 43-46*) Purified human platelet GPIbα ectodomain was digested with trypsin. Tryptic peptides were purified and then treated with peptide-N-glycosidase F (PNGase F) to release N-glycans. Released N-glycans were purified by liquid chromatography. O-glycans were released from N-glycan-free GPIbα glycopeptides by reductive β-elimination. Further details are available in the Supporting Information.

### 2.5 MS glycomics analysis

Released N- and O-glycans were analyzed as their permethylated forms by nanospray ionization mass spectrometry (NSI-MS) in positive mode.(*40, 43-46*) Permethylated glycans were dissolved in 50 µL of 1 mM sodium hydroxide in methanol/water (1:1) for infusion into an Orbitrap linear ion trap mass spectrometer (Orbitrap-LTQ; Thermo Fisher Scientific) using a nanospray source at a syringe flow rate of 0.80 µL/min and capillary temperature of 210 °C. For fragmentation by collision-induced dissociation (CID) in MS/MS and MS^n^, a normalized collision energy of 35 – 40% was used. MS data were manually annotated using the Xcalibur software package version 2.0 (Thermo Fisher Scientific) as previously described.(*40, 47*) Glycans were also analyzed by total ion mapping (TIM) to detect minor glycan components. Graphic representations of N-glycan monosaccharide residues were consistent with the Symbol Nomenclature for Glycans (SNFG) as adopted by the glycomics and glycobiology communities.(*48, 49*) The angle at which branching monosaccharide residues are depicted in graphic representations is not meant to infer specific linkage positions or anomeric configuration, some of which remain to be determined by future analysis. Glycomics data and metadata were obtained and are presented in accordance with MIRAGE standards and the Athens Guidelines.(*50*) All raw mass spectrometry data were deposited at GlycoPost, accession #GPST000275.(*51*) Further details are available in the Supporting Information.

### 2.6 In vitro digestion of GPIbα ectodomain for glycoproteomics

The ectodomain of GPIbα was digested with several different conditions to maximize protein coverage and glycopeptide identification. When proteases were included in the digestion, peptides were subsequently reduced in 2 mM dithiothreitol (DTT) at 65 °C for 30 minutes. After cooling, iodoacetamide (IAA) was added to a concentration of 3 mM and allowed to react for 15 minutes in the dark at room temperature. Samples were then diluted using 50 µL of 50 mM ammonium bicarbonate. All digests were performed in ammonium bicarbonate, pH 7.5. Mucinase digests (e.g., StcE, SmE) were performed overnight and protease (e.g., GluC, trypsin) digests were performed the following day for 6 h. In cases where N-glycans were removed, PNGaseF (NEB) was diluted 1:10 and 1 µL was added to the mucinase digests (20-60 µL) overnight. In sialidase treated samples (recombinant protein only), 1 µL of sialoEXO (Genovis) was added to 39 µL of water, then 1 µL of the dilution was added to the reaction. All digests were incubated at 37 °C. The combinations of proteases used and their enzyme : substrate ratios were as follows: SmE/GluC (1:20, 1:100), SmE/trypsin (1:20, 1:100), StcE/GluC (1:20, 1:100), StcE/Thermolysin (1:20, 1:100), SmE only (1:20), ImpA only (1:20). Reactions were quenched using 100 µL of 0.5% formic acid in ultrapure water (Pierce). C18 clean-up was performed using 1 mL strataX columns (Phenomenex). Each column was hydrated with 1 mL of acetonitrile, followed by one 1-mL rinse of 0.1% formic acid in water (“buffer A”). The samples were then added to the column and rinsed with 150 µL of 0.1% formic acid. The samples were eluted twice with 150 µL of 0.1% formic acid in 30% acetonitrile and dried by vacuum centrifugation. Finally, a HILIC clean-up was performed to remove polyethylene glycol (PEG) contamination. The samples were reconstituted in 10 µL of buffer A for MS analysis. Further details are available in the Supporting Information.

### 2.7 Glycoproteomics MS data acquisition and analysis

Samples were analyzed by online nanoflow liquid chromatography (LC)-tandem MS using an Orbitrap Eclipse Tribrid mass spectrometer (Thermo Fisher Scientific) coupled to a Dionex Ultimate 3000 HPLC (Thermo Fisher Scientific). Higher-energy collisional dissociation (HCD), electron transfer dissociation, and electron transfer dissociation with supplemental activation (EThcD) were performed. Raw files were searched using O-Pair search with MetaMorpheus and Byonic (ProteinMetrics) against a database containing the protein of interest.(*52, 53*) Files were searched with various cleavage specificities dependent on digestion conditions and search algorithm used. For O-Pair, an O-glycan database was built based on the glycomics data generated in this study and the maximum number of glycosites per peptide was set to 4. For Byonic, the O-Glycan common 9 database was used with a common max of 3. Peptide hits were filtered using a 1% FDR and, in O-Pair, a q value of < 0.01. All peptides were manually validated and/or sequenced using Xcalibur software (Thermo Fisher Scientific). Candidate sequences were obtained from search algorithms and an in-house program was used to identify MS2 spectra containing oxonium ions and the associated naked peptide backbone. HCD spectra were used to assign naked peptide sequences, glycan compositions, and characteristic fragmentation patterns. ETD and/or EThcD were then used to localize glycosites. The mass spectrometry proteomics data have been deposited to the ProteomeXchange Consortium via the PRIDE partner repository with the dataset identifier PXD035030.(*54, 55*) Further details are available in the Supporting Information.

## 3. Results

### 3.1 Western Blot Analysis of Human Platelet GPIbα Ectodomain Confirms Presence of ABH Antigens

Given our interest in the potential hemostatic relevance of GPIbα ABH antigens, we first aimed to validate prior reports that this protein bears ABH antigens. We purified the GPIbα ectodomain (glycocalicin) from human apheresis platelets of blood type A, B, or O (Figure S3-S4, Table S4), and subjected the purified protein to Western blot analysis with anti-A or anti-B antibodies (Figure 2).

**Figure 2.**
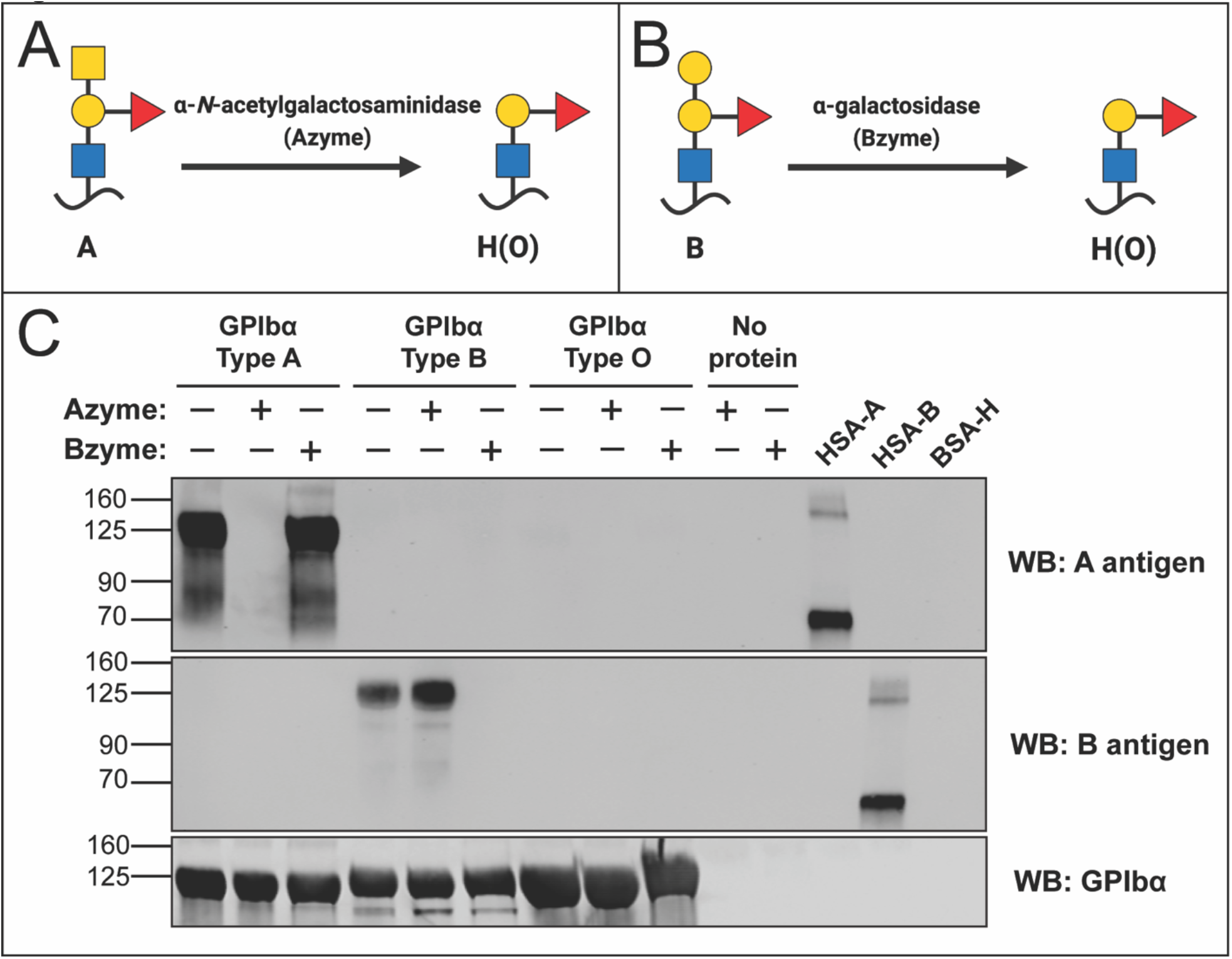
GPIbα carries A and B antigens. A) α-*N*-acetyl-galactosaminidase (“Azyme”) hydrolysis of the terminal GalNAc of the A antigen to yield the H antigen. B) α-galactosidase (“Bzyme”) hydrolysis of the terminal Gal of the B antigen to yield the H antigen. C) GPIbα ectodomain purified from human platelets of blood type A, B, or O was untreated, or treated with Azyme or Bzyme. The resulting proteins were analyzed by Western blot with anti-A and anti-B antibodies. The following synthetic glycoproteins were used as controls: human serum albumin covalently modified with A antigen (HSA-A), human serum albumin covalently modified with B antigen (HSA-B), and bovine serum albumin covalently modified with the H antigen (BSA-H). Blue square, *N*-acetylglucosamine (GlcNAc); yellow circle, Galactose (Gal); red triangle, L-Fucose (Fuc).

Type A GPIbα, but not B or O, showed reactivity with anti-A antibody, and this reactivity was abolished when the protein was treated with a recombinant α-*N*-acetylgalactosaminidase that cleaves the terminal GalNAc residue found in the A antigen (Figure 2A, 2C). Similarly, type B GPIbα, but not A or O, showed reactivity on Western blot with anti-B antibody, and this reactivity was abolished with recombinant α-galactosidase treatment which cleaves the terminal Gal residue found in the B antigen (Figure 2B, 2C).

To test for H antigen, we performed lectin blot analysis using *Ulex europaeus* (UEA-I) lectin. UEA-I lectin binds fucose-containing glycans and is selective for the H antigen over A or B. GPIbα from type A, B, and O platelets showed reactivity on UEA-I lectin blot (Figure 3). Individuals with non-O blood types are known to express H antigen, so UEA-I reactivity on the protein from type A and B platelets was not surprising.(*38, 56*) As expected, no UEA-I reactivity was seen following fucosidase treatment. As further validation of the presence of the H antigen on GPIbα from type O platelets, type O GPIbα was treated with recombinant human ABO glycosyltransferase type B. This led to a band that showed reactivity on an anti-B Western blot (Figure S5). Enzymatic removal of N-glycans from A or B GPIbα with PNGaseF did not substantially diminish the A or B signal on Western blot, suggesting that ABH antigens do not reside exclusively on N-glycans (Figure S6-S8).

**Figure 3.**
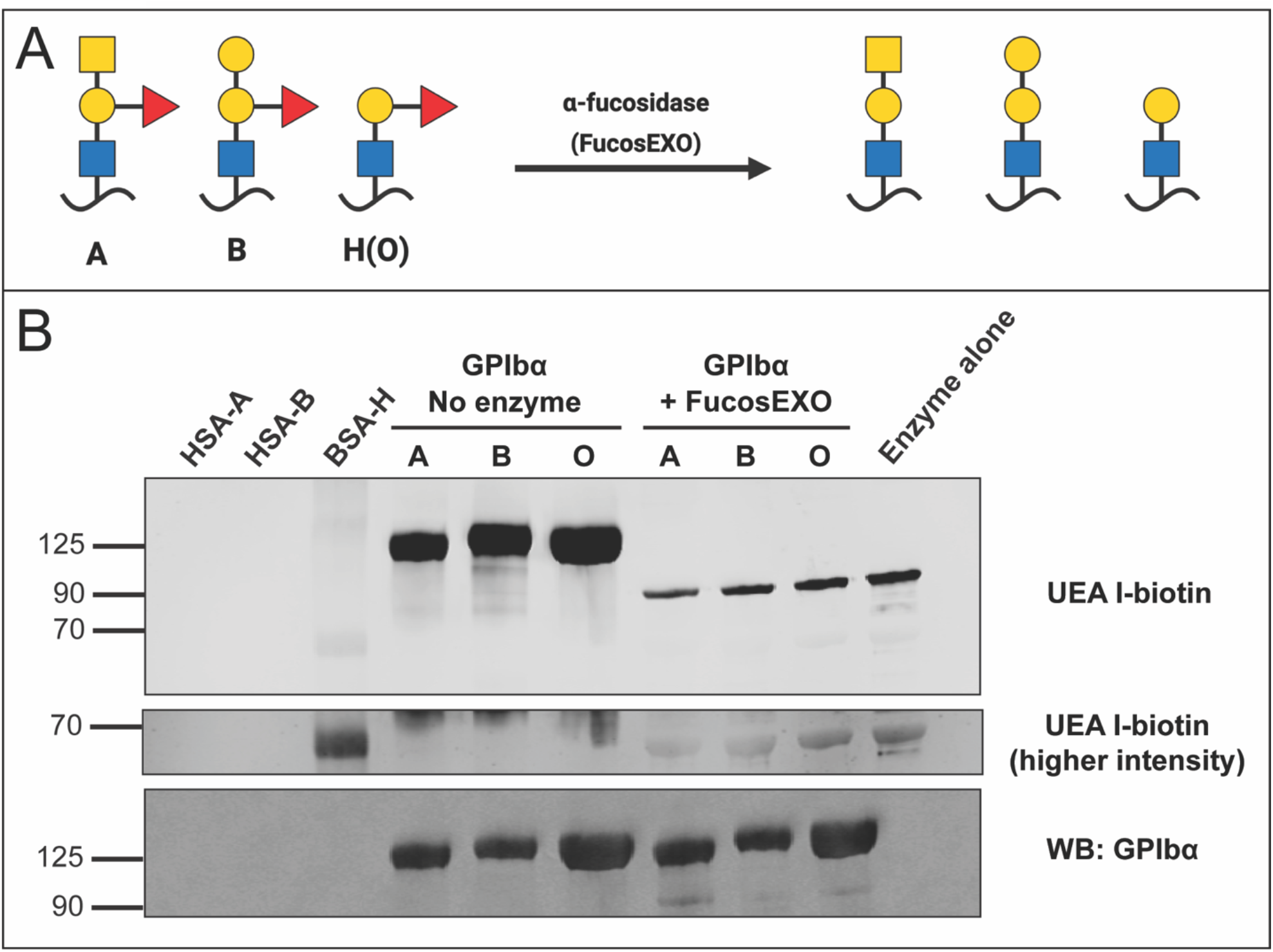
GPIbα carries the H antigen. A) α-fucosidase (FucosEXO) hydrolysis of ABH antigen fucose to yield defucosylated antigens. B) GPIbα ectodomain purified from human platelets of blood type A, B, or O was untreated or treated with fucosEXO, and then analyzed by blot with *Ulex europaeus* agglutinin I (UEA-I). UEA-I binds to α-fucosylated glycans, and preferentially binds to H antigens. The following synthetic glycoproteins were used as controls: human serum albumin covalently modified with A antigen (HSA-A), human serum albumin covalently modified with B antigen (HSA-B), and bovine serum albumin covalently modified with the H antigen (BSA-H). Blue square, *N*-acetylglucosamine (GlcNAc); yellow circle, Galactose (Gal); red triangle, L-Fucose (Fuc).

### 3.2 Quantification of Human Platelet GPIbα Ectodomain Monosaccharides Confirms Extensive Glycosylation

We next sought to confirm that GPIbα is heavily glycosylated and to quantify monosaccharides of interest. We were particularly interested in determining the abundance of 1) SA and 2) fucose, which is a constituent of ABH antigens. Purified GPIbα ectodomain from blood type O human apheresis platelets was treated with acid to hydrolyze the glycans into monosaccharides. The monosaccharides were quantified by established liquid chromatography methods (Table S5-S6). The most common type of SA in humans is *N*-acetylneuraminic acid (Neu5Ac). This analysis revealed the following molar ratios of monosaccharide to protein (mean +/-standard dev): Neu5Ac 58.6 +/-0.1, Gal 56.5 +/-3.2, GlcNAc 33.2 +/-0.7, GalNAc 21.1 +/-0.6, Man 6.7 +/-0.1, Fuc 5.1 +/-0.2, glucose 2.0 +/-0.3 (Figure S9-10, Table S7).

### 3.3 MS Glycomics Analysis of Human Platelet GPIbα Ectodomain Reveals a Diverse Repertoire of N- and O-glycans

After validating that GPIbα carries ABH antigens and SA, we aimed to determine what glycans bear these moieties by MS glycomics analysis. Purified GPIbα ectodomain from blood type A, B, or O human apheresis platelets was subjected to trypsin digestion and PNGaseF treatment to release N-glycans. The N-glycans were saved and de-N-glycosylated glycopeptides were subjected to *β*-elimination to release O-glycans. Separately, N- and O-glycans were permethylated and analyzed by MS. The glycan sequences for isobaric structures were determined by MS^n^ fragmentation.

Eight distinct O-glycans were identified that were > 3% relative abundance on the full MS profile (Figure 4, Figure S11). The most abundant O-glycan in each sample was a disialylated core 2 hexasaccharide (*m/z* 864.4, 1705.8); which accounted for approximately 30-50% of the total O-glycan ion current in each sample. Most of the other observed O-glycans were core 1 or core 2 type O-glycans with variable monosaccharide extensions. Seven of the O-glycans were sialoglycans, carrying 1 to 3 SA monomers per glycan. Tn and T antigen were not able to be accurately detected or quantified by glycomics given their low masses.

**Figure 4.**
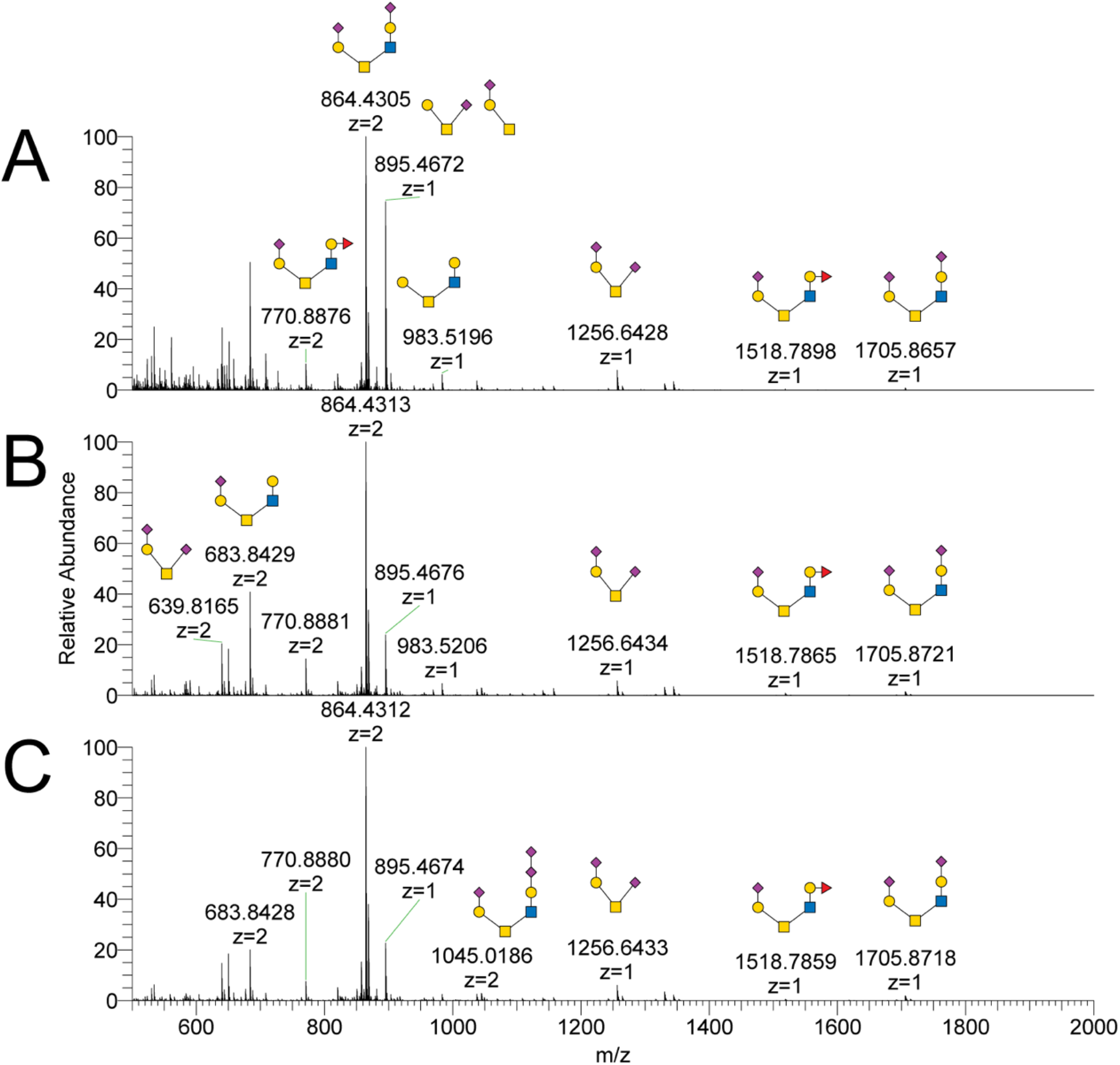
GPIbα O-glycan profiling by MS. O-glycans were released from GPIbα ectodomain purified from human platelets of blood type A, B, or O, respectively. The data shown here are raw MS profiles that include multiply charged ions. GPIbα purified from platelets of blood type: A) A, B) B, and C) O. The structures of isobaric O-glycans were characterized by MS^n^ analysis. Blue square, *N*-acetylglucosamine (GlcNAc); green circle, Mannose (Man); yellow circle, Galactose (Gal); purple diamond, *N*-acetylneuraminic acid (Neu5Ac); red triangle, L-Fucose (Fuc).

A terminally fucosylated complex core 2 O-glycan (*m/z* 770.9, 1518.8) was detected in samples from blood types A, B, and O. The relative abundance was approximately 5% in the type O sample. MS^n^ analysis of this glycan confirmed the sequence as expected for the H antigen (Figure 5). The MS^3^ fragment ions for the putative H antigen glycans were those expected based on fragmentation of an H antigen standard; these were distinct from fragment ions seen with a Lewis antigen standard (Figure S12). Three other O-glycans were found to bear the H antigen. These were < 3% relative abundance in the full MS spectrum and were identified by TIM analysis and confirmed by MS^n^ (Figure S13). These low-abundance H-antigen O-glycans include structures where the H-antigen is a terminal modification of complex core 2 O-glycans as well as mucin type H antigen, where the H antigen is a direct modification of Ser/Thr (Figure 1D).

**Figure 5.**
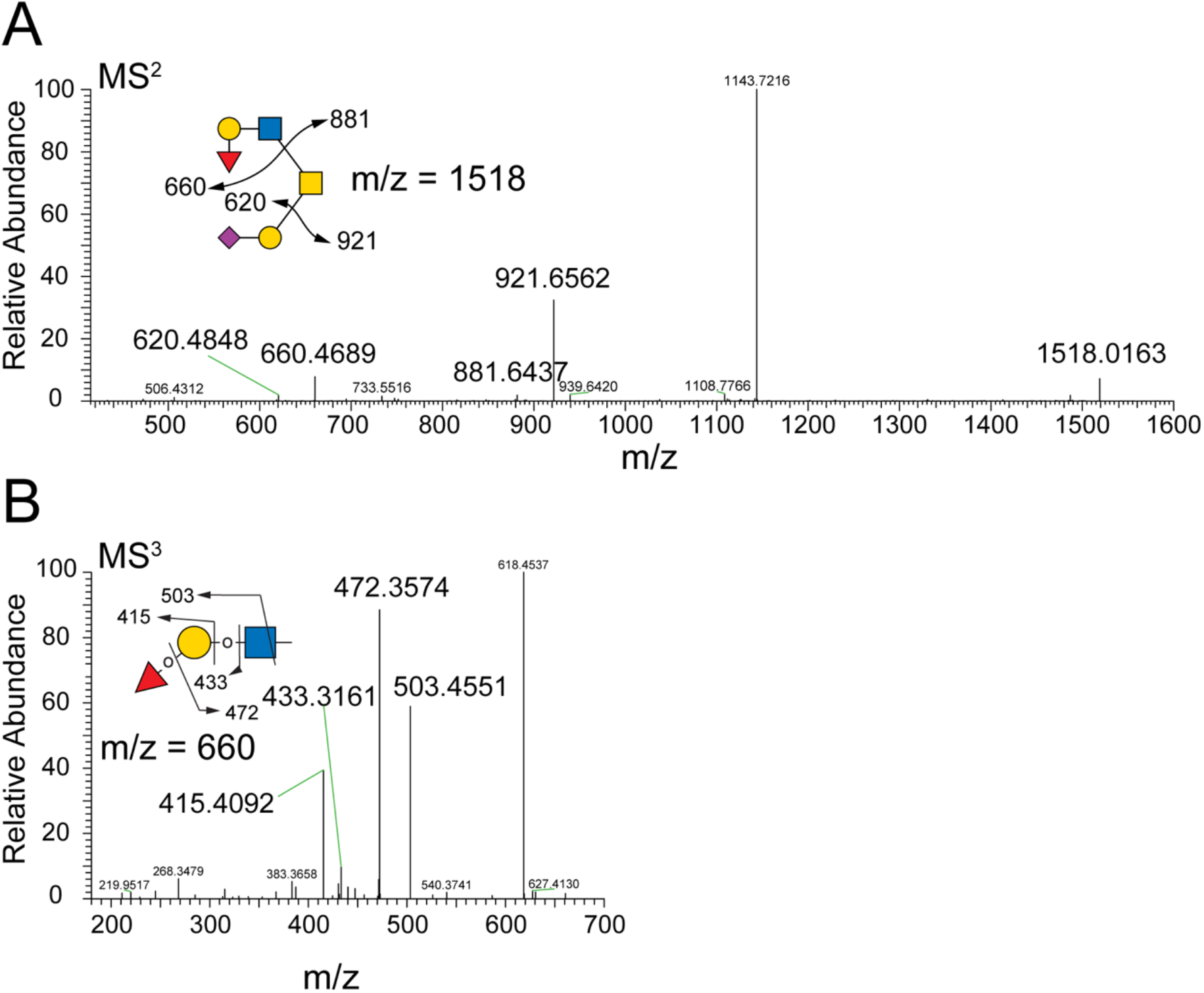
Determination of H antigen on a core 2 O-glycan by MS^n^. O-glycomics analysis of type O GPIbα revealed a glycan with m/z 1518; this is compatible with an extended core 2 O-glycan carrying the H antigen. A) MS^2^ fragmentation of the m/z 1518 O-glycan generated a fragment at m/z 660, compatible with the trisaccharide that defines the H antigen. B) MS^3^ fragmentation of the m/z 660 fragment generated masses consistent with the H antigen (m/z = 503, 472, 433, 415). Blue square, *N*-acetylglucosamine (GlcNAc); yellow circle, Galactose (Gal); purple diamond, *N*-acetylneuraminic acid (Neu5Ac); red triangle, L-Fucose (Fuc).

In analysis of type A GPIbα, two O-glycans bearing A antigen epitopes were detectable in the intact MS profile, but the signal was minimal and these glycans were not confirmed by MS^n^. The B antigen was observed on three O-glycans in the analysis of type B GPIbα and could be confirmed by MS^n^ analysis (Figure S13-S14). H antigen was also observed on O-glycans from GPIbα from blood type A and B platelets, consistent with the positive signal seen on UEA-I lectin blots for GPIbα of all blood types.

Eighteen distinct N-glycan masses were identified that were > 3% relative abundance in the full MS profile (Figure 6, Figure S15). The most abundant N-glycans in each sample were complex bi-antennary disialylated glycans (*m/z* 2792.5, *m/z* 2966.5). All identified N-glycans were of the complex type except for one high Man N-glycan (*m/z* 1579.8). The complex N-glycans had up to five antennae. Seventeen distinct N-glycans were identified that contain SA, and there were up to four SA monomers per glycan.

**Figure 6.**
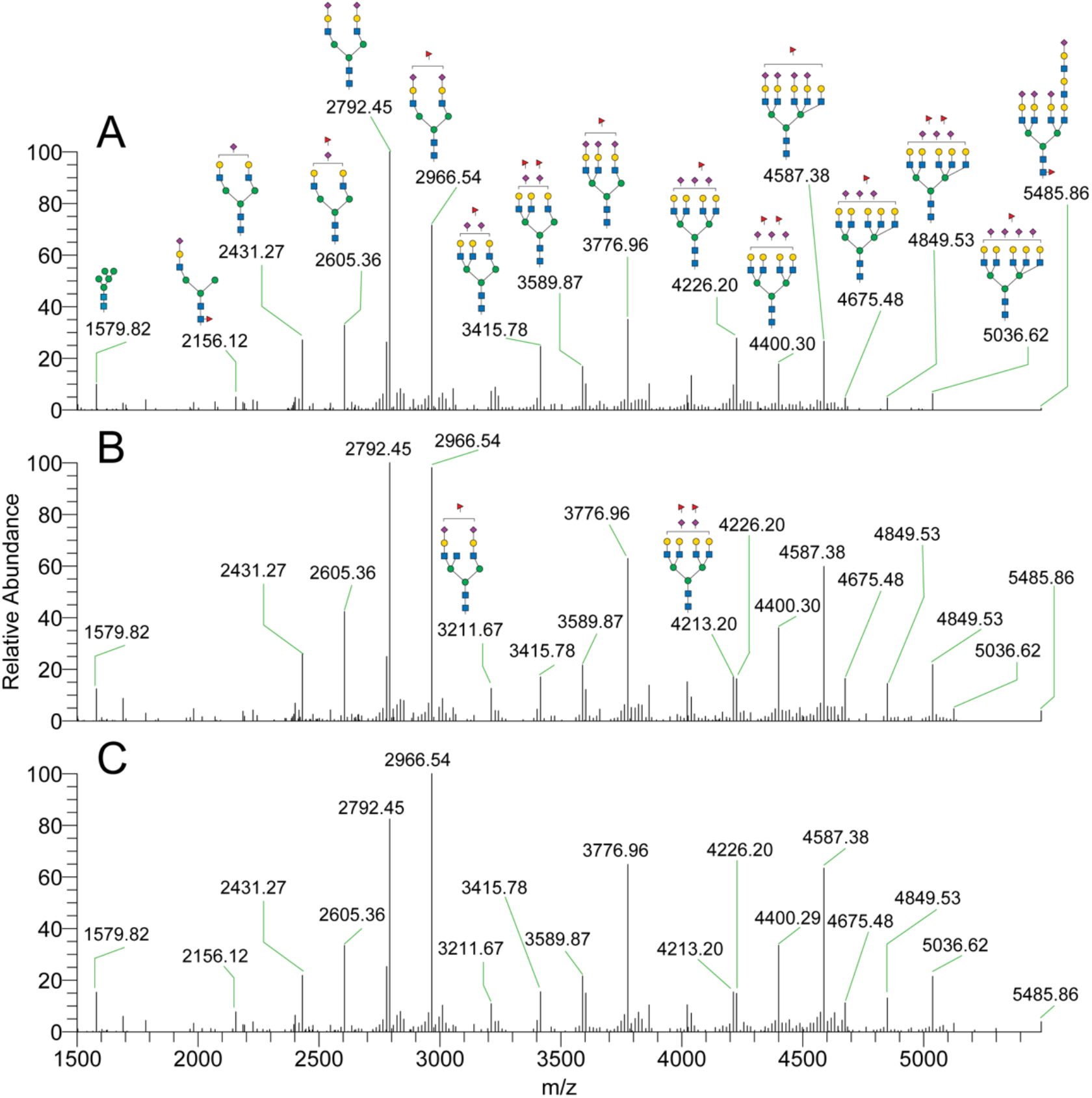
GPIbα N-glycan profiling by MS. N-glycans were released from GPIbα ectodomain purified from human platelets of blood type A, B, or O, respectively. Raw MS profiles were deconvoluted by Xtract software version 2.0. GPIbα purified from platelets of blood type: A) A, B) B, and C) O. The structures of isobaric N-glycans were characterized by MS^n^ analysis. Blue square, *N*-acetylglucosamine (GlcNAc); green circle, Mannose (Man); yellow circle, Galactose (Gal); purple diamond, *N*-acetylneuraminic acid (Neu5Ac); red triangle, L-Fucose (Fuc).

N-glycomics analysis of type O GPIbα revealed 10 N-glycans with masses compatible with an H antigen epitope, which was confirmed by MS^n^ fragmentation (Figure S16-17). The A antigen was detected on two N-glycans from type A GPIbα and was confirmed by MS^n^ analysis (Figure S17-18). No B antigen was detected on N-glycans released from type B GPIbα. H antigen was also observed on N-glycans from GPIbα from blood type A and B platelets (Figure S17).

### 3.4 Bioinformatic Analysis of GPIbα Ectodomain Predicts 62 Glycosites

We used bioinformatic analysis of the GPIbα amino acid sequence to generate hypotheses regarding the identity of the glycosites. N-glycosites are predicted by the consensus amino acid motif for N-glycosylation: N-X-S/T, where X can be any amino acid except proline.(*57*) Analysis of the GPIbα ectodomain amino acid sequence reveals four putative N-glycosites, including two in the VWF LBD (N37, N175) and one in the mucin domain (N398) (Figure 7, Table S2).(*57*) There is no consensus motif for O-glycosylation. The NetOGlyc 4.0 server produces neural network predictions of O-glycosites based on previously characterized O-glycoproteins.(*58*) NetOGlyc 4.0 predicted 58 GPIbα O-glycosites, including two in the VWF LBD (T256 and S257) and 25 in the mucin domain (Figure 7, Table S2).

**Figure 7.**
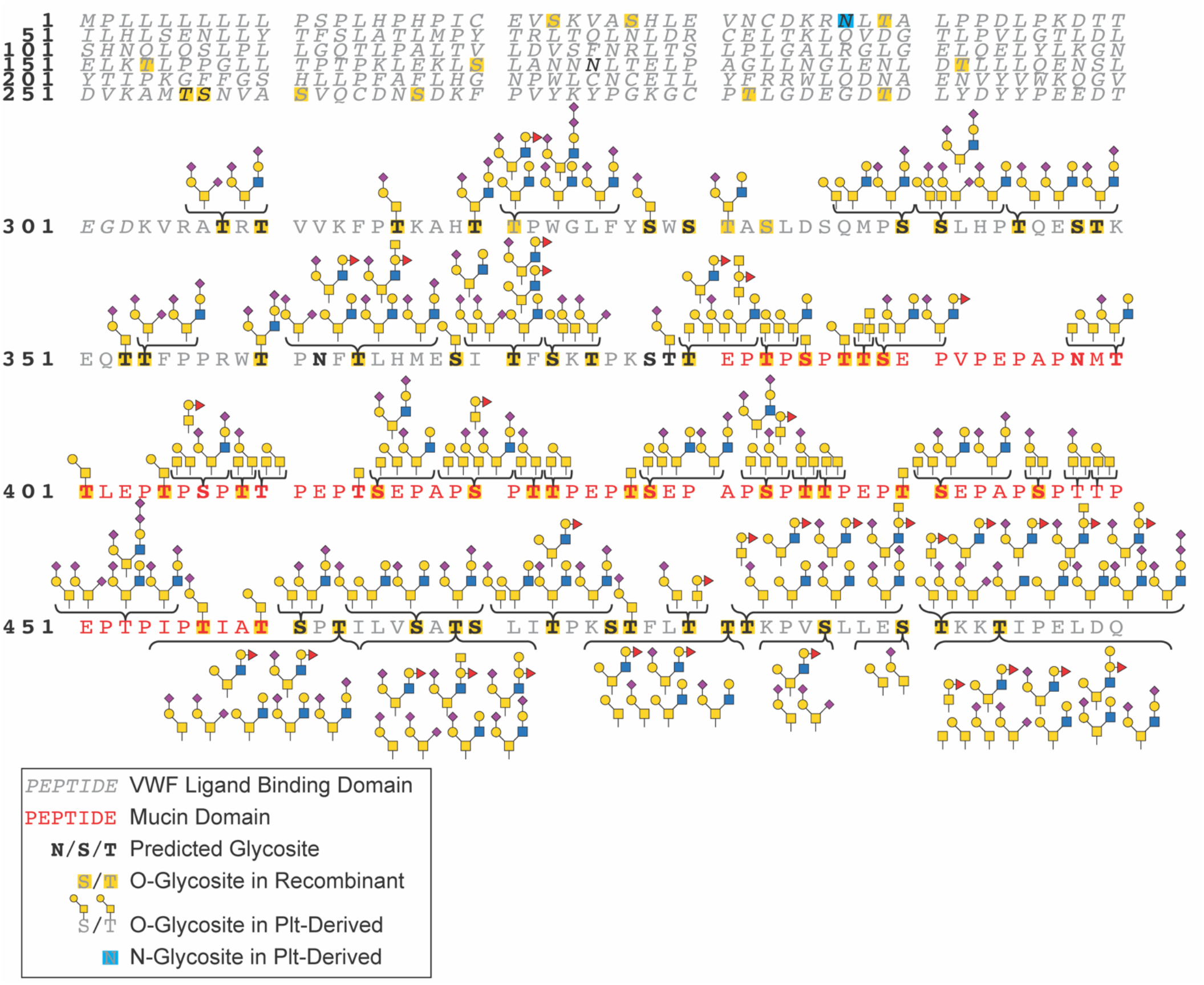
GPIbα glycosites and glycan structures determined in this analysis. The GPIbα ectodomain amino acid sequence here is from UniprotKB – P07359.2. The Von Willebrand Factor ligand binding domain residues are in italics and mucin domain residues are red. Bolded amino acids are computationally predicted glycosites by NetGlyc. O-glycosites identified in the analysis of the recombinant protein are highlighted yellow. The N-glycosite identified in the analysis of the protein purified from human platelets highlighted teal. O-glycosites identified in the analysis of the protein purified from human platelets are indicated with glycans listed above or below the amino acid. The UniprotKB – P07359.2 amino acid sequence includes 3 tandem repeats. The amino acid sequence is identical for three stretches of amino acids: 407-419, 420-432, and 433-445. Identified glycosites within these repeats are listed as being present in all 3 repeats, but the sites could not be unambiguously assigned to a particular repeat. VWF, Von Willebrand Factor; Plt, platelet; Blue square, *N*-acetylglucosamine (GlcNAc); green circle, yellow circle, Galactose (Gal); purple diamond, *N*-acetylneuraminic acid (Neu5Ac); red triangle, L-Fucose (Fuc).

### 3.5 Glycoproteomics Analysis of GPIbα Glycosites

We next aimed to map GPIbα glycosites experimentally by glycoproteomics. We first studied recombinant human GPIbα ectodomain by subjecting GPIbα to digestion with various combinations of bacterial mucinases (StcE and SmE), glycosidases, and commercially available proteases. We analyzed O-glycosites and O-glycans from the resultant glycopeptides by LC-MS/MS. We identified 62 unique O-glycosites in the recombinant protein, including 16 O-glycosites in the mucin domain (Figure 7, Table S2). One or more O-glycan masses was identified at each of the glycosites (Table S8).

We used similar methods to perform a glycoproteomic analysis of platelet-derived GPIbα ectodomain purified from human platelets, this time seeking to characterize both O- and N-glycosites. We identified 48 unique O-glycosites and 1 N-glycosite in the platelet-derived protein, including 21 unique O-glycosites within the mucin domain. GPIbα from biological samples has variable numbers of tandem repeats (1-4). If we consider GPIbα with 3 tandem repeats (as in the standard Uniprot sequence), and we assume that each repeat has the same glycosites, then there would be a total of 60 glycosites in the ectodomain and 32 in the mucin domain. In the VWF LBD, we identified 1 N-glycosite (N37).

The Tn antigen was identified at 14 glycosites, the T antigen at 19 glycosites (Table S2). With knowledge of the structures of the SA and ABH antigen-bearing glycans from the glycomics analysis, we used the glycoproteomics data to site-localize these glycans. We identified 38 O-glycosites which bear sialoglycans and 15 O-glycosites which bear ABH antigens (Figure 7, Table S2).

## 4. Discussion

Here we report the most detailed analysis of platelet GPIbα glycosylation to date; we have experimentally determined 49 unique glycosites and 26 major glycan structures by MS.

Our monosaccharide analysis validates prior reports which suggested that GPIbα is heavily glycosylated.(*9-12*) Our data confirm that the GPIbα ectodomain carries abundant SA, with a monosaccharide to protein molar ratio of 59 for Neu5Ac. The approximately 2:2:1:1 ratio of Neu5Ac:Gal:GalNAc:GlcNAc is consistent with monosaccharide analyses performed in some of the earliest biochemical characterizations of the GPIbα ectodomain.(*9*) Furthermore, the monosaccharide analysis provides insight into the relative ratios of N-vs O-glycans. Each N-glycan typically contains at least three Man units, while Man is characteristically absent from mucin-type O-glycans.(*13*) Therefore, the monosaccharide to protein molar ratio of 59 for Neu5Ac, 57 for Gal, and only 7 for Man is consistent with a preponderance of O-glycans compared with N-glycans. Finally, the detection of Fuc supports the presence of fucose-containing glycans, of which ABH antigens are one subtype.

Of the four potential GPIbα N-glycosites, two have been previously reported to be occupied by N-glycans (N37, N175).(*29, 31*)We confirmed occupancy at N37. We did not detect N-glycosylation at any other N-glycosites. Of the 58 bioinformatically predicted GPIbα O-glycosites, 8 have been previously experimentally determined to be occupied by O-glycans.(*29, 30*) In our analysis of the protein purified from platelets, we reconfirmed occupancy at 7 out of 8 of these glycosites; we did not observe O-glycosylation at the previously determined S470. Two VWF ligand-binding domain O-glycosites were predicted bioinformatically (S256 and S257); these were observed to be glycosylated in our analysis of the recombinant protein but not the platelet-derived protein. As anticipated from prior work and bioinformatic analysis of the amino acid sequence, our glycoproteomic analysis revealed that the GPIbα mucin domain is densely O-glycosylated. We identified 21 O-glycosites in the GPIbα mucin domain. O-glycosites were also found in the broader macroglycopeptide domain and MSD.

Given the critical role of SA in regulating platelet clearance, it is notable that 1) SA (Neu5Ac) was the most abundant monosaccharide in the GPIbα compositional analysis, 2) we detected 24 abundant glycans that terminated in SA, and 3) we identified 39 glycosites that were occupied by sialoglycans. The mechanism by which desialylated platelets are cleared remains incompletely understood. Some studies have suggested that the mechanism involves exposure of GPIbα Gal residues, followed by platelet uptake via interactions with hepatic and/or Kupffer cell asialoglycoprotein receptors.(*59*) Recent reports point towards a role for desialylation of GPIbα O-glycans.(*7*) Our site-localization of GPIbα sialoglycans provides a map of potential glycans that could be relevant in mediating platelet clearance.

Numerous epidemiologic studies have demonstrated that risk of bleeding and thrombosis are associated with ABO blood type.(*27, 60-63*) Blood type O individuals are at higher risk of bleeding and lower risk of thrombosis compared with non-O individuals. Lower levels of VWF and factor VIII (FVIII) in blood type O compared with non-O individuals are likely important drivers of this association.(*64*) However, two recent studies have led to the consideration that there may be a platelet-intrinsic mechanism that also underlies this association.(*27, 63*) First, in a genome-wide association study, the *ABO* locus was found to be the main determinant of primary hemostasis as measured by Platelet Function Analyzer (PFA-100) closure times.(*65*) The influence of ABO blood type on PFA-100 measurements was only partially explained by VWF and FVIII levels, suggesting that other mechanisms may be important.(*65*) Second, in a microfluidics assay, non-O platelets were found to have superior binding characteristics to VWF compared with type O platelets.(*26*) For the first time, we have definitively demonstrated the presence of ABH antigens on GPIbα glycans by robust Western blot assay and MS. We identified 15 glycosites which bear ABH antigen-containing glycans, largely located in the mucin domain and MSD. Whether GPIbα ABH antigen glycans affect the structure and/or function of this protein, potentially leading to changes in VWF binding, remains to be tested in future work.

Deletion of a stretch of 9 amino acids in the MSD (462-470, PTILVSATS) causes inherited platelet-type VWD.(*66*) We confirmed glycosylation of the protein from platelets at 3 glycosites within this sequence (T463, S467, T469) and identified ABH antigen glycans at 2 of these sites (T463 and T469). A fourth glycosite within this stretch, S470, has been previously characterized.(*30*) Further studies are required to determine the relevance of glycosylation at these 4 glycosites to VWD.

The Tn and T antigens are biosynthetic precursors to more fully elaborated O-glycan structures. We identified these immature O-glycans commonly in more densely O-glycosylated stretches of GPIbα. For example, the mucin glycosites T385, T387, and T388 carried only Tn, T, or GalNAc-GalNAc, and not more mature O-glycans. This preponderance of immature O-glycan structures in densely glycosylated regions has been observed in analysis of other proteins and may relate to steric hindrance of glycosyltransferase activity in nascent, heavily O-glycosylated proteins.(*67*)

Anti-platelet glycoprotein antibodies lead to accelerated platelet clearance in several immune thrombocytopenia syndromes: ITP, fetal-neonatal alloimmune thrombocytopenia (FNAIT), post-transfusion purpura (PTP), and the Tn syndrome.(*8, 23, 25, 68, 69*) Prior work suggests that these antibodies may bind to platelets in a glycan-dependent manner.(*23, 25, 69*) For example, in a recent study, anti-T antigen antibodies were identified in sera from a cohort of pediatric ITP patients and implicated in the pathogenesis of ITP.(*23*) Thrombocytopenia can develop in the Tn syndrome when anti-Tn antibodies bind to platelets. We have now established that GPIbα carries T and Tn antigen epitopes. This opens the door to investigation into the relevance of auto-antibodies directed against GPIbα T or Tn antigen in the pathogenesis of immune thrombocytopenia syndromes. More broadly, our characterization of GPIbα glycosylation provides insight into possible glycoepitopes that may be targeted by anti-glycan and/or anti-GPIb allo- or auto-antibodies in immune thrombocytopenia syndromes.

Some limitations of our work should be noted. We studied the glycosylation of GPIbα ectodomain that was cleaved from the surface of human platelets. The glycosylation of the cleaved ectodomain may not be the same as the glycosylation of the intact protein on the platelet surface. O-glycosylation is known to regulate proteolytic cleavage of many proteins, so certain glycoforms of GPIbα could be preferentially proteolyzed by ADAM17.(*70*) Furthermore, we purified the protein using lectin and anion exchange chromatography; these purification steps could have selectively enriched certain glycoforms. While our analysis has revealed many more glycosites and glycans than had previously been characterized, we may not have identified all the glycosites and glycans for two major reasons: 1) dense and heterogeneous glycosylation may render any individual glycopeptide too low abundance to detect and/or might hinder ionization of the glycopeptide, 2) we did not search for glycopeptide masses compatible with glycosylation and other post-translational modifications to the same peptide, so we would not detect glycosites located near non-glycan modifications. While we performed glycomics analysis on preparations of GPIbα that were pure enough to yield a single band on Coomassie-stained SDS-PAGE, it is possible that some of the glycans observed by MS glycomics came from contaminating proteins.

## 5. Conclusion

In summary, we report a comprehensive analysis of GPIbα glycosylation. We used MS glycomics and glycoproteomics to perform a site-specific analysis of GPIbα purified from human platelets. We determined 48 O-glycosites and 1 N-glycosite. We determined that GPIbα carries diverse N- and O-glycans, including sialoglycans, Tn antigen, T antigen, and ABH antigens. This information lays the foundation for further studies to determine the functional and structural implications of GPIbα glycosylation in hemostasis, platelet clearance, and the anti-platelet immune response.

## Acknowledgements

Monosaccharide analysis was performed by the UC San Diego GlycoAnalytics Core. This work was supported by a Sarafan ChEM-H Physician-Scientist fellowship (to MAH), the Stanford Maternal & Child Health Research Institute Instructor K Award Support Program (to MAH), a National Blood Foundation Early Career Scientific Research Grant (to MAH), a NIH NHBLI Pathway to Independence award (1K99HL156029-01 to MAH), and NIH grant R01CA200423 (to CRB). KA was supported by the GaLSIC collaborative research fund, Soka University, Japan. SAM is supported by the Yale Science Development Fund and the Yale SEAS/Science Program to Advance Research Collaboration (SPARC). KEM is supported by a Yale Endowed Postdoctoral Fellowship and SCL is supported by the National Institutes of Health Chemical Biology Training Grant (T32 GM067543). Some figures were created with BioRender.com.

## Authorship Details

Marie Hollenhorst, Carolyn Bertozzi, and Stacy Malaker designed the project. Marie Hollenhorst, Katherine Tiemeyer, Keira Mahoney, Kazuhiro Aoki, Mayumi Ishihara, Valentina Rangel-Angarita, Sarah Lowery, and Stacy Malaker performed the experiments. Marie Hollenhorst and Katherine Tiemeyer wrote the manuscript. All authors interpreted data and revised the manuscript.

## Footnotes

1 GPIbα amino acids are numbered throughout the manuscript according to the UniprotKB – P07359.2 (GP1BA_HUMAN). The GPIbα gene is polymorphic; it codes for GPIbα protein with a variable number (1-4) of tandem 13-amino acid repeats (SEPAPSPTTPEPT). This sequence has 3 tandem repeats. See Figure S1 for more details regarding the amino acid sequence.

