## Supporting Information for "Comprehensive analysis of platelet glycoprotein Ibα glycosylation"

Table of Contents — Page

**I. Supplemental Materials and Methods** — **S4-11**

**A.** Key resources table. — S4

**B.** Apheresis platelet donor ABO blood grouping. — S5

**C.** GPIbα ectodomain purification from human platelets. — S6

**D.** Calculation of GPIbα concentration from 280 nm absorbance. — S6

**E.** Western blotting for detection of GPIbα, ABH antigens, and sialic acid. — S7

**F.** GalNAcEXO (α-*N*-acetylgalactosaminidase) treatment of GPIbα. — S8

**G.** α-galactosidase treatment of GPIbα. — S8

**H.** PNGase F release of GPIbα N-glycans for Western blotting. — S8

**I.** Fucosidase treatment of GPIbα. — S8

**J.** Conversion of H antigen to B antigen using glycosyltransferase B. — S8

**K.** Quantification of GPIbα sialic acid by DMB assay. — S8

**L.** Quantification of GPIbα monosaccharides by anion exchange chromatography. — S9

**M.** Release of GPIbα N-glycans for glycomics analysis. — S9

**N.** Release of O-glycans from GPIbα N-glycan-free-glycopeptides for glycomics analysis. S10

**O.** Glycoproteomics MS data acquisition and analysis. S10

**II. Supplemental Figures S12-29**

**Figure S1.** GPIbα amino acid sequence. S12

**Figure S2.** Glycans determined in prior analyses of GPIbα. S13

**Figure S3.** Anion exchange chromatography purification of GPIbα ectodomain. S14

**Figure S4.** SDS-PAGE and anti-GPIbα blot analysis of GPIbα purification. S15

**Figure S5.** Glycosyltransferase B converts GPIbα H antigens into B antigens. S16

**Figure S6.** PNGase F treatment of type A GPIbα does not eliminate A antigens. S17

**Figure S7.** PNGase F treatment of type B GPIbα does not eliminate B antigens. S18

**Figure S8.** PNGase F de-N-glycosylates fetuin. S19

**Figure S9.** Sialic acid quantification by DMB assay. S20

**Figure S10.** Monosaccharide analysis by high-performance anion-exchange chromatography with pulsed amperometric detection (HPAEC-PAD). S21

**Figure S11.** O-glycans identified in glycomics analysis of GPIbα. S22

**Figure S12.** $MS^3$ fragmentation of a Lewis antigen standard. S23

**Figure S13.** O-glycans that carry ABH antigens identified in glycomics analysis of GPIbα. S24

**Figure S14.** Type B GPIbα carries B antigen on core 2 type O-glycan. S25

**Figure S15.** N-glycans identified in glycomics analysis of GPIbα. S26

**Figure S16.** Type O GPIbα carries H antigen on a tetra-antennary complex N-glycan. S27

**Figure S17.** N-glycans that carry ABH antigens identified in glycomics analysis of GPIbα. S28

**Figure S18.** Type A GPIbα carries both A and H antigen on bi-antennary complex N-glycans. S29

**III. Supplemental Tables S30-37**

**Table S1.** Domain organization of the GPIbα ectodomain. S30

**Table S2.** GPIbα ectodomain glycosites and glycan structures determined previously and in this analysis. S30

**Table S3.** Platelet donor ABO blood group determination. S32

**Table S4.** Representative GPIbα ectodomain purification yields. S33

**Table S5.** Gradient used for UPLC separation of DMB-sialic acid. S33

**Table S6.** Gradient used for HPAEC-PAD separation of monosaccharides. S33

**Table S7.** Monosaccharide composition of GPIbα ectodomain. S34

**Table S8.** Glycan structures identified in glycoproteomics analysis of recombinant GPIbα ectodomain. S35

**IV. Supplemental References S37-38**

### I. Supplemental Materials and Methods

#### A. Key resources table.

| REAGENT or RESOURCE | SOURCE | IDENTIFIER | NOTES |
|---|---|---|---|
| Antibodies and Lectins | | | |
| Rabbit monoclonal anti-GPIbα/CD42b (clone EPR19204) | Abcam | Cat# ab210407; RRID: AB_2909590 | WB 1:10,000 |
| Ortho Clinical Diagnostics ABO Blood Group Reagents: Anti-A | Thermo Fisher Scientific | Cat# 23-287-451 Ref#210520 RRID: AB_2909591 | WB 1:10 |
| Ortho Clinical Diagnostics ABO Blood Group Reagents: anti-B | Thermo Fisher Scientific | Cat# 23-287-451 Ref#210521 RRID: AB_2909591 | WB 1:10 |
| Ulex Europaeus Agglutinin I (UEA I), biotinylated | Vector Laboratories | SKU B-1065-2 | WB 5 µg/mL |
| Sambucus Nigra Lectin (SNA, EBL), biotinylated | Vector Laboratories | SKU B-1035-2 | WB 2 µg/µL |
| IRDye 800 CW goat anti-mouse IgM | LI-COR | P/N 926-32280 RRID: AB_2814919 | WB 1:15,000 |
| IRDye 800 CW goat anti-rabbit | LI-COR | P/N 926-32211 RRID: AB_621843 | WB 1:15,000 |
| IRDye 800 CW Streptavidin | LI-COR | P/N 926-32230 | WB 1:1000 |
| Biological samples | | | |
| Expired human apheresis platelets of blood type A, B, or O | Bloodworks Northwest | 2965-02 Expired apheresis platelets | Up to 7 days out of date |
| Recombinant proteins | | | |
| Recombinant human CD42b/GPIbα protein, CF | R&D Systems | Cat# 4067-GP-050 | Recombinant human protein, expressed in an NS0-derived mouse myeloma cell line |
| Blood Group A-human serum albumin (6-atom spacer) | Dextra Laboratories | Product code NGP9305 | |
| Blood Group B- human serum albumin (6-atom spacer) | Dextra Laboratories | Product code NGP9323 | |
| Blood Group H disaccharide-bovine serum albumin (3-atom spacer) | Dextra Laboratories | Product code NGP0205 | |
| GalNAcEXO | Genovis | G1-NA1-020 | Recombinant *Akkermansia muciniphila* exo-α-*N*-acetylgalactosaminidase, expressed in *E. coli* |
| Recombinant human blood group B transferase | R&D Systems | Cat# 6824-GT-020 | Recombinant human enzyme, expressed in a Chinese Hamster Ovary cell line |
| α-Galactosidase from green coffee beans | Sigma Aldrich | CAS 9025-35-8; SKU G8507 | |
| FucosEXO | Genovis | G1-FM1-020 | Two His-tagged fucosidases recombinantly expressed in *E. coli* |
| SialEXO | Genovis | G1-SM1-020 | Mix of sialidases derived from *Akkermansia muciniphila*, recombinantly expressed *in E. coli* |

| | | | |
|---|---|---|---|
| Peptide-N-Glycosidase F (for Western blot analysis) | New England BioLabs | Cat# P0704S | Peptide-N-Glycosidase F (for Western blot analysis) |
| Peptide-N-Glycosidase F (for N-glycomics analysis) | Prozyme | Code GKE-5006 | Gene from *Chryseobacterium meningosepticum* expressed in *E. coli* |
| Endoproteinase Glu-C (GluC) | Roche | CAS: 137010-42-5 | From *Staphylococcus aureus* |
| O-glycoprotease (ImpA) | NEB | Cat# P0761S | |
| Thermolysin | Promega | Part# V400A | |
| Secreted protease of C1 esterase inhibitor (StcE) | | | *E. coli* enzyme, purified as previously described.(*1*) |
| Protein Purification and Analysis | | | |
| XT MOPS buffer | Bio-Rad | Cat# 1610788 | |
| Intercept PBS Blocking Buffer | LI-COR | P/N: 927-90001 | |
| Wheat Germ Agglutinin (WGA), agarose bound | Vector Laboratories | AL-1023 | |
| HiTrap DEAE Sepharose Fast Flow, 1 mL | Cytiva Life Sciences | P/N 17505501 | |
| Zeba Spin Desalting Columns | Thermo Fisher Scientific | Cat# 89882, 89889, 89891, 89893 | 0.5 – 10 mL resin bed volume, 7-40K MWCO |
| Amicon Ultra-15 centrifugal filter units | Millipore Sigma | Cat# UCF910024 | 100 KDa MWCO |
| Chameleon Duo Prestained Protein Ladder | LI-COR | P/N 92860000 | |
| 4-12% Criterion XT Precast Bis-Tris gel | Bio-Rad | Cat# 3450123, 3450124, 3450125 | |
| Criterion Gel Electrophoresis Cell | Bio-Rad | Cat# 1656001 | |
| AcquaStain Protein Gel Stain | Bulldog Bio | Cat# AS001000 | |
| Trans-Blot Turbo | Bio-Rad | Cat# 1704150 | |
| Carbo-Free Blocking Solution (10X concentrate) | Vector Laboratories | SKU SP-5040-125 | |
| Revert 700 Total Protein Stain | LI-COR | P/N 926-11011 | |
| NewBlot Nitro Stripping Buffer for Nitrocellulose Membranes | LI-COR | P/N 928-40030 | |
| Deposited data | | | |
| MS Glycomics | GlycoPost https://glycopost.glycosmos.org/ | #GPST000275 | |
| MS Glycoproteomics | PRIDE https://www.ebi.ac.uk/pride/ | ID PXD035030 | **Username:** **Password:** FOwHUl4s |
| Gel and blot full images | Mendeley Data, V1 https://data.mendeley.com/ | DOI:10.17632/6fw3gg5v5x.1 | |

#### B. Apheresis platelet donor ABO blood grouping.

Donor ABO blood group was determined at Bloodworks Northwest according to standard operating procedure for donor ABO blood group determination. Testing was performed using a Beckman Coulter PK3700 Automated Microplate System. A forward type was determined by incubating donor red blood cells (RBCs) with anti-A, anti-B, or anti-A,B antibodies and assessing for agglutination. A reverse type was determined by incubating donor plasma with type A1 or B test RBCs. ABO blood type was assigned based on the results of this test according to Table S3.

**C. GPIbα ectodomain purification from human platelets.**

The soluble ectodomain of GPIbα was purified from out of date apheresis platelets using an adaptation of previously reported protocols.(*2, 3*) Apheresis platelet units of known blood type were obtained from BloodWorks Northwest. Platelets were transferred to 500 mL conical centrifuge tubes and pelleted for 40 min at 2,000 x g at room temperature. The supernatant was discarded, and platelet pellets were resuspended in an equivalent volume of ice-cold phosphate-buffered saline containing no calcium or magnesium (PBS). From this point on, the platelet suspension was maintained at 4 °C. Platelet suspensions were re-centrifuged at 2000 x g for 40 min at 4 °C. The supernatant was discarded, and each pellet was resuspended in approximately 10 mL of cold 10 mM Tris-HCl pH 7.4 containing 150 mM NaCl and 2 mM $CaCl_2$. Platelet suspensions were transferred to 50 mL conical tubes and subjected to sonication with a Misonix Sonicator 3000 Ultrasonic Cell disruptor for two 15-second intervals on setting 4, separated by a 60 second period of rest on ice. Following sonication, platelets were incubated at 37 °C for 1 hour. After incubation, platelet suspensions were centrifuged for 30 min at 10,000 x g at 4 °C.

The supernatant was loaded onto a lectin wheat germ agglutinin (WGA) agarose (Vector Laboratories) gravity column. Approximately 100 mg of protein was loaded per mL of WGA agarose. Prior to loading, the column was washed with 5 column volumes of 50 mM Tris-HCl, pH 7.4 containing 10 mM $MgCl_2$ (wash buffer). The platelet lysate was loaded onto the column and allowed to flow through by gravity or with gentle, manual positive pressure. The flow-through was re-loaded onto the column and allowed to flow through a second time. The column was washed twice with 5 column volumes of wash buffer. Glycoproteins were eluted in fractions of 0.2 column volumes using wash buffer containing 0.25 M *N*-acetylglucosamine, prepared fresh the day of the purification.

GPIbα-containing fractions were concentrated by centrifugation in a 100 kDa molecular weight cut-off (MWCO) centrifugal filter unit (Amicon). The buffer was then exchanged using a spin desalting column (Zeba) to 20 mM Tris-HCl pH 7.4. The protein was further purified by anion exchange chromatography using a HiTrap DEAE-FF 1 mL column (Cytiva) on an AKTA pure liquid chromatography system (Cytiva). Buffer A was 20 mM Tris-HCl, pH 7.4, and buffer B was 20 mM Tris-Cl with 0.7 M NaCl. The flow rate was 0.5 mL/min. The column was washed with 6 column volumes of 100% buffer A, then protein was eluted using a gradient from 15% to 60% B over 20 column volumes (Figure S3). GPIbα-containing fractions of highest purity generally eluted between 16-22 column volumes, as determined by immunoblot and total protein stain. These fractions were pooled, concentrated by MWCO centrifugal filter unit, and buffer exchanged into 50 mM ammonium bicarbonate, pH 7.5 (Table S4). Aliquots of protein solution were stored at -80 °C.

**D. Calculation of GPIbα concentration from 280 nm absorbance.**

The molar extinction coefficient ($\sum_{molar}$) of GPIbα amino acids 1-505 (ectodomain) was determined to be 51,225 $M^{-1}$ $cm^{-1}$ using the Expasy ProtParam tool under the assumption that all cysteines are reduced.(*4*) The percent extinction coefficient ($\sum$1%) was determined as follows:

$$\varepsilon 1\% = \frac{(\varepsilon_{molar})(10)}{Molecular\ weight} = \frac{(51225)(10)}{55401.25} = 0.925$$

Absorbance at 280 nm was measured in triplicate using a NanoDrop 2000 spectrophotometer (Thermo Scientific) with the setting of 1 absorbance unit = 1 mg/mL. The average of these values was then multiplied by the percent extinction coefficient for GPIbα ($\sum 1\%$ = 0.925) to give total protein concentration of the sample.

**E. Western blotting for detection of GPIbα, ABH antigens, and sialic acid.**

Purified GPIbα (5 μg) was separated by sodium dodecyl sulfate-polyacrylamide gel electrophoresis (SDS-PAGE) using a 4-12% Criterion XT Precast Bis-Tris gel (Bio-Rad) in 1x XT MOPS buffer (Bio-Rad). The protein was transferred to a 0.45 um nitrocellulose membrane for immunoblotting using a Trans-Blot Turbo (Bio-Rad) transfer system. Membranes were blocked in Intercept (PBS) blocking buffer (Li-COR) overnight at 4 °C. Blots were incubated with primary antibody diluted in Intercept PBS blocking buffer containing 0.2% Tween-20 for 1 hour at room temperature. The following primary antibodies were used: anti-A or anti-B murine monoclonal blend (1:10, Ortho-Clinical Diagnostics), rabbit monoclonal anti-GPIbα (1:10,000, Abcam ab 210407, EPR19204). Primary antibody incubation was followed by 4 x 5 min washes in phosphate buffered saline (PBS) with 0.1% Tween-20. Secondary antibodies were diluted in Intercept PBS blocking buffer containing 0.2% Tween-20 (for GPIbα and lectin blotting) or 0.4% Tween-20 (for anti-A/B blotting). Blots were incubated with the following secondary antibodies for 1 hour at room temperature: IRDye 800 CW goat anti-mouse IgM (LiCOR, 1:15,000), IRDye 800 CW goat anti-rabbit (Li-COR, 1:15,000). Secondary antibody incubation was followed by 4 x 5 min washes in PBS-T (0.1% Tween-20 for GPIbα and lectin blotting, 0.2% Tween-20 for anti-A/B blotting). Blots were rinsed briefly in PBS before imaging on the Odyssey DLx imaging system (Li-COR).

H antigen was visualized using biotinylated *Ulex europaeus* I (UEA-I, Vector Laboratories). Blots were blocked at 4 °C overnight in 1X Carbo-Free blocking solution (Vector Laboratories), followed by incubation with UEA-I (biotin) at a concentration of 5 μg/mL in 1X Carbo-Free blocking buffer containing 0.1% Tween-20. Biotinylated lectin incubation was followed by 4 x 5min washes in PBS-T (0.1% Tween-20) and then a 1 hour, room temperature incubation in secondary antibody (IRDye 800 CW Streptavidin, 1:1000 in Carbo-Free blocking buffer containing 0.2% Tween-20). Blot was washed 4 x 5min in PBS-T (0.1% Tween-20) and imaged as described above. Sialic acid-containing GPIbα glycans were also assessed by lectin blotting using the *Sambucus nigra* (SNA/EBL) biotinylated lectin (Vector Laboratories) at a working concentration of 2 μg/μL. For SNA lectin blotting, 0.1 mM $CaCl_2$ was added to the Carbo-free blocking buffer during lectin incubation.

Commercial synthetic glycoproteins bearing ABH antigens were used as controls: blood group A-HSA, blood group B-HSA, and blood group H-BSA (Dextra Laboratories).

To re-probe blots, antibody was stripped from the membrane using 1X-5X New Blot Nitro stripping buffer for nitrocellulose membranes (LI-COR). Membranes were incubated with stripping buffer for 5-15 min, rinsed 3 times with PBS, and imaged to ensure all signal was removed. Blots were blocked again before re-probing.

**F. GalNAcEXO (α-*N*-acetylgalactosaminidase) treatment of GPIbα.**

Purified GPIbα was treated with GalNAcEXO, an α-*N*-acetylgalactosaminidase (Genovis). 1 unit of GalNAcEXO was added per µg of glycoprotein. Reactions were brought up to a final volume of 20-30 ∝L using 20mM Tris-HCl, pH 7.4 and incubated for 2 hours at 37 °C.

**G. α-galactosidase treatment of GPIbα.**

Purified GPIbα was treated with α-galactosidase from green coffee beans (Sigma-Aldrich), which is supplied as an ammonium sulfate suspension. The suspension was centrifuged at 10,000 x g for 10 min at 4 °C. The ammonium sulfate supernatant was discarded, and the pelleted enzyme was resuspended in an equal volume of 50 mM sodium phosphate buffer, pH 6.5, to reach a final concentration of 45.6 units/mL. Enzyme was added to a final reaction concentration of 32 mU per ∝g of GPIbα, in a final concentration of 50 mM sodium phosphate buffer, pH 6.5. Final reaction volumes ranged from 10 – 63 ∝L, and reactions were incubated at 37 °C overnight.

**H. PNGase F release of GPIbα N-glycans for Western blotting.**

PNGase F digestion was carried out according to manufacturer's protocol under denaturing conditions (New England BioLabs). First, GPIbα at a concentration of 1.2 µg/µL in 1X PNGase F reaction buffer in a total volume of 5 µL was heated at 95 °C for 30 minutes to denature the protein. This solution was cooled on ice, and then 1 µL of 10X glycobuffer, 1 µL of 10% NP-40, and 3 µL of water were added. Next, 1.5 µL of PNGase F was added to bring the final volume to 11.5 µL. The reaction was incubated at 37 °C for 60 minutes. Enzymatic activity of PNGase F was confirmed by de-N-glycosylation of fetuin monitored by molecular weight shift on SDS-PAGE (Figure S8).

**I. Fucosidase treatment of GPIbα.**

Purified GPIbα from all blood types was treated with FucosEXO (Genovis). Final reaction conditions were as follows: 0.67 units FucosEXO per µL (8U FucosEXO/µg of GPIbα), 0.5 µg/µL GPIbα, 20 mM Tris-HCl pH 6.8 in a final reaction volume of 12 µL. Reactions were incubated overnight at 37 °C.

**J. Conversion of H antigen to B antigen using glycosyltransferase B.**

GPIbα from type O individuals was treated with recombinant human blood group B transferase (R&D Systems). Reaction conditions were adapted from manufacturer's recommendation as follows: 12.5 µg/mL blood group B transferase (4.77 ng enzyme per µg of GPIbα), 0.26 µg/µL GPIbα (5.5 µg total), 1.5 mM UDP-galactose, 5 mM $MnCl_2$, and 50 mM HEPES pH 6.5 in a total reaction volume of 21 µL. Reactions were incubated at 37 °C for 16 hours, and antigen conversion was confirmed by western blotting.

**K. Quantification of GPIbα sialic acid by DMB assay.**

Quantification of sialic acid was performed using an adaptation of previously reported methods.(*5-7*) GPIbα (10 ng) was treated with 2 M acetic acid at 80 °C for 3 h followed by removal of excess acid using a speed vacuum. Sialic acid was then tagged using 1,2-diamino-4,5-methyleneoxybenzene (DMB). The DMB reagent mixture was prepared fresh before reaction. To prepare sufficient reagent for tagging 5 samples, 0.87

mg of DMB was dissolved in 162.3 µL of Milli-Q water, followed by the addition of 44.3 µL glacial acetic acid, 29.1 µL 2-mercaptoethanol and 39.6 µL of (0.25 M) sodium hydrosulfite and mixed thoroughly. Samples were dissolved in 50 µL water and 50 µL freshly prepared DMB reagent mixture was added per sample. The DMB reaction was carried out at 50ºC for 2.5 hours. Reactions were allowed to cool before being analyzed by reversed phase ultra performance liquid chromatography, fluorescence (RP-UPLC-FL, Waters Acquity). Samples were eluted with a mixture of Solvent A (7% methanol in water) and Solvent B (100% acetonitrile) over 18 minutes. A Bridge Ethylene Hybrid (BEH) C18 column (2.1 × 50 mm, Waters) was used at a flow rate of 0.4 mL/min for profiling DMB-sialic acid; the gradient details are in Table S5. The excitation and emission wavelengths were 373 nm and 448 nm, respectively. Known amounts of standard Neu5Ac and Neu5Gc were used to quantify sialic acids in the samples.

**L. Quantification of GPIbα monosaccharides by anion exchange chromatography.**

Quantification of monosaccharides was performed using an adaptation of previously reported methods.(*8, 9*) 4.2 µg of GPIbα was treated with 100 µL of 2 M trifluoroacetic acid at 100 °C for 4 hr followed by removal of excess acid by dry nitrogen flush. To ensure complete removal of acid, the samples were co-evaporated twice with isopropanol at a 1:1 ratio. Samples were then dissolved in ultra-pure distilled water and injected on Dionex ICS3000 (Thermo Scientific) using a CarboPac PA-1 column (4 mm x 250 mm) attached to a Carbo PA1-guard column (4 mm x 50 mm). Data were acquired using the standard Quad waveform for carbohydrates supplied by the manufacturer of the Dionex ICS3000. The solvent gradient is shown in Table S6; Solvent A was HPLC grade water, solvent B was 100 mM NaOH, 7 mM sodium acetate and solvent C was 100 mM NaOH, 25 mM NaOAc. The flow rate was 1.0 mL/min and the total run time 50 min. Monosaccharides were quantified by comparison with the peak area of known amount of standard mixture of L-fucose, D-galactosamine, D-glucosamine, D-galactose, D-glucose, and D-mannose.

**M. Release of GPIbα N-glycans for glycomics analysis.**

N-glycans were released from 70-100 µg of GPIbα ectodomain purified from human platelets as previously reported.(*10-12*) GPIbα was resuspended in 200 µL of trypsin buffer (0.1 M Tris-HCl, pH 8.2, containing 1 mM $CaCl_2$) by sonication and boiled for 5 min. After cooling to room temperature, 25 µL of trypsin solution (2 mg/mL in trypsin buffer) was added. Digestion was allowed to proceed for 18 h at 37 °C before the mixture was boiled for 5 min. The reaction mixture was dried by vacuum centrifugation. The dried peptide and glycopeptide mixture was resuspended in 200 µL of 5% acetic acid (v/v) and loaded onto a Sep-Pak C18 cartridge column. The cartridge was washed with 5 column volumes of 5% acetic acid. Glycopeptides were eluted stepwise, first with 3 volumes of 20% isopropyl alcohol in 5% acetic acid and then with 3 volumes of 40% isopropyl alcohol in 5% acetic acid. The 20 and 40% isopropyl alcohol steps were pooled and evaporated to dryness. Dried glycopeptides were resuspended in 50 µl of 50 mM sodium phosphate buffer, pH 7.5, for digestion with PNGaseF. Following PNGase digestion for 18 h at 37 °C, released N-linked glycans were separated from peptides and enzyme by passage through a Sep-Pak C18 cartridge. The digestion mixture was reconstituted in 200 µL of 5% acetic acid and loaded onto a Sep-Pak C18 cartridge column. The column passthrough and an additional elution with 3 volumes of 5% acetic acid, containing released N-glycans, were collected combined, and evaporated to dryness. O-glycopeptides were recovered from the C18 Sep-Pak cartridge column by eluting with 3 volumes of 100% isopropyl alcohol.

**N. Release of O-glycans from GPIbα N-glycan-free-glycopeptides for glycomics analysis.**

O-linked glycans were released from N-glycan free glycopeptides by β-elimination under reductive conditions.(*12-14*) (*12-14*) O-glycopeptides were reconstituted in 250 µL of 100 mM sodium hydroxide containing 1 M sodium borohydride and incubated for 18 hrs at 45 °C in a glass tube sealed with a Teflon-lined screw top. The tube was transferred to ice and acetic acid added to 10% to neutralize the reaction. The sample was then loaded onto a column of DOWEX 50W X 8 cation exchange resin (Bio-Rad) to desalt. Released O-glycans were eluted from the resin with three bed-volumes of 5% acetic acid and lyophilized to dryness. To remove borate from the sample, a solution of 10% acetic acid in methanol was added and the sample was then dried under a stream of nitrogen gas at 37 °C; this was repeated for a total of five times. The sample was then resuspended in 5% acetic acid and loaded onto a C18 Sep-Pak cartridge column (Waters) that was previously washed with acetonitrile and pre-equilibrated with 5% acetic acid. Run-through from the column was collected after loading; the column was then washed a total of five times with 5% acetic acid. The run-through and washes were combined and evaporated to dryness.

**O. Glycoproteomics MS data acquisition and analysis.**

Samples were analyzed by online nanoflow liquid chromatography-tandem mass spectrometry using an Orbitrap Eclipse Tribrid mass spectrometer (Thermo Fisher Scientific) coupled to a Dionex Ultimate 3000 HPLC (Thermo Fisher Scientific). Mucinase/protease digests (0.5-1 µg) or mucinase only digests (1-5 µg) were loaded via autosampler isocratically onto a C18 nano precolumn using buffer A. For preconcentration and desalting, the column was washed with 2% acetonitrile and 0.1% formic acid in water (“loading pump solvent”). Subsequently, the C18 nano precolumn was switched in line with the C18 nano separation column (75-µm × 250-mm EASYSpray containing 2 µm C18 beads) for gradient elution. The column was held at 35 °C using a column heater in the EASY-Spray ionization source (Thermo Fisher Scientific). The samples were eluted at a constant flow rate of 0.3 µL/min using a 60-min gradient. The gradient profile was as follows: 0-0-35-95-95-2% B in 0-5-65-70-75-77 minutes, respectively.

The instrument method used an MS1 resolution of 60,000 full width at half maximum (FWHM) at 400 m/z, an automatic gain control (AGC) target of 3e5, and a mass range from 300 to 1,500 m/z. Dynamic exclusion was enabled with a repeat count of 3, repeat duration of 10 s, and exclusion duration of 10 s. Only charge states 2 to 6 were selected for fragmentation. MS2s were generated at top speed for 3 s. Higher-energy collisional dissociation (HCD) was performed on all selected precursor masses with the following parameters: isolation window of 2 m/z, 28% collision energy, orbitrap detection (resolution of 7,500), max inject time of 75 ms, and an AGC target of 1e4 ions. Electron-transfer dissociation with supplemental activation (EThcD) was performed if 1) the precursor mass was between 300 and 1,500 m/z and 2) 3 of 9 HexNAc or NeuAc fingerprint ions (126.055, 138.055, 144.07, 168.065, 186.076, 204.086, 274.092, and 292.103) were present at ± 0.1 m/z and greater than 5% relative intensity. EThcD parameters were as follows: Orbitrap detection (resolution 7,500) calibrated charge-dependent ETD times, 15% nce for HCD, maximum inject time of 250 ms, reagent AGC target of 5e5, and precursor AGC target of 1e5. In some instances, samples were re-injected and subjected to an HCD-pd-ETD method where HCD and product-dependent triggering were the same as above, but ETD parameters were as follows: ion trap detection, calibrated charge-dependent ETD times, maximum inject time of 100 ms, reagent AGC target of 5e5, and precursor AGC target of 5e4.

Raw files were searched using O-Pair search with MetaMorpheus and Byonic (ProteinMetrics) against a database containing the protein of interest. Files were searched with various cleavage specificities. In all searches, N-terminal S/T was selected; in tryptic digests, C-terminal K/R; in GluC digests, C-terminal E; 10 missed cleavages were allowed. Mass tolerance was set at 10 ppm for MS1s and 20 ppm for MS2s. Cysteine carbamidomethylation was set as a fixed modification and methionine oxidation was allowed as a variable modification. For O-Pair, an O-glycan database was built based on the glycomics data generated in this study and the maximum number of glycosites per peptide was set to 4. For Byonic, the O-Glycan common 9 database was used with a common max of 3. Peptide hits were filtered using a 1% FDR and, in O-Pair, a q value of < 0.01. All peptides were manually validated and/or sequenced using Xcalibur software (Thermo Fisher Scientific).

Candidate sequences were obtained from search algorithms and an in-house program was used to identify MS2 spectra containing oxonium ions and the associated naked peptide backbone. HCD spectra were used to assign glycan compositions and characteristic fragmentation patterns. ETD and/or EThcD were then used to localize glycosites.

#### II. Supplemental Figures

**Figure S1. GPIbα amino acid sequence.**

Recombinant GPIbα ectodomain was purchased from R & D Biosystems. This is the human protein recombinantly expressed in a mouse NS0-derived cell line. The amino acid sequence for the protein is based on the UniprotKB/Swiss-Prot P07359.1.(*15*) The recombinant protein encompasses His17 to Leu505 from the P07359.1 sequence, with a C-terminal 6-His tag. This amino acid sequence differs from the updated UniprotKB/Swiss-Prot P07359.2. The P07359.2 sequence has an additional 26 amino acids starting at amino acid 428: SEPAPSPTTPEPTSEPAPSPTTPEPT. Throughout the manuscript, amino acid numbers are assigned based on the UniprotKB/Swiss-Prot P07359.2 amino acid numbering.

Amino acid sequence of recombinant GPIbα used in this study (based on UniprotKB/Swiss-Prot P07359.1).

```
  1                 hpic evskvashle vncdkrnlta lppdlpkdtt ilhlsenlly
 61 tfslatlmpy trltqlnldr celtklqvdg tlpvlgtldl shnqlqslpl lgqtlpaltv
121 ldvsfnrlts lplgalrglg elqelylkgn elktlppgll tptpklekls lannnltelp
181 agllnglenl dtlllqensl ytipkgffgs hllpfaflhg npwlcnceil yfrrwlqdna
241 envyvwkqgv dvkamtsnva svqcdnsdkf pvykypgkgc ptlgdegdtd lydyypeedt
301 egdkvratrt vvkfptkaht tpwglfysws tasldsqmps slhptqestk eqttfpprwt
361 pnftlhmesi tfsktpkstt eptpspttse pvpepapnmt tleptpsptt peptsepaps
421 pttpeptpip tiatsptilv satslitpks tflttttkpvs llestkktip eldqppklrg
481 vlqghlessr ndpflhpdfc cllplhhhhh h
```

Amino acid sequence of GPIbα UniprotKB/Swiss-Prot P07359.2. Amino acid numbering used in this manuscript is based on this sequence. The additional 26 amino acids are bolded.

<pre>
  1 mplllllllll psplhphpic evskvashle vncdkrnlta lppdlpkdtt ilhlsenlly
 61 tfslatlmpy trltqlnldr celtklqvdg tlpvlgtldl shnqlqslpl lgqtlpaltv
121 ldvsfnrlts lplgalrglg elqelylkgn elktlppgll tptpklekls lannnltelp
181 agllnglenl dtlllqensl ytipkgffgs hllpfaflhg npwlcnceil yfrrwlqdna
241 envyvwkqgv dvkamtsnva svqcdnsdkf pvykypgkgc ptlgdegdtd lydyypeedt
301 egdkvratrt vvkfptkaht tpwglfysws tasldsqmps slhptqestk eqttfpprwt
361 pnftlhmesi tfsktpkstt eptpspttse pvpepapnmt tleptpsptt peptsepaps
421 pttpeptsep apspttpept sepapspttp eptpiptiat sptilvsats litpkstflt
481 ttkpvslles tkktipeldq ppklrgvlqg hlessrndpf lhpdfccllp lgfyvlglfw
541 llfasvvlil llswvghvkp qaldsgqgaa lttatqtthl elqrgrqvtv prawllflrg
601 slptfrsslf lwvrpngrvg plvagrrpsa lsqgrgqdll stvsirysgh sl
</pre>

**Figure S2. Glycans determined in prior analyses of GPIbα.** Glycans with gray boxes are structures that are compatible with H antigen-bearing glycans. Of note, these structures were not described as H antigen glycans in the papers reporting their characterization.(*16-21*)

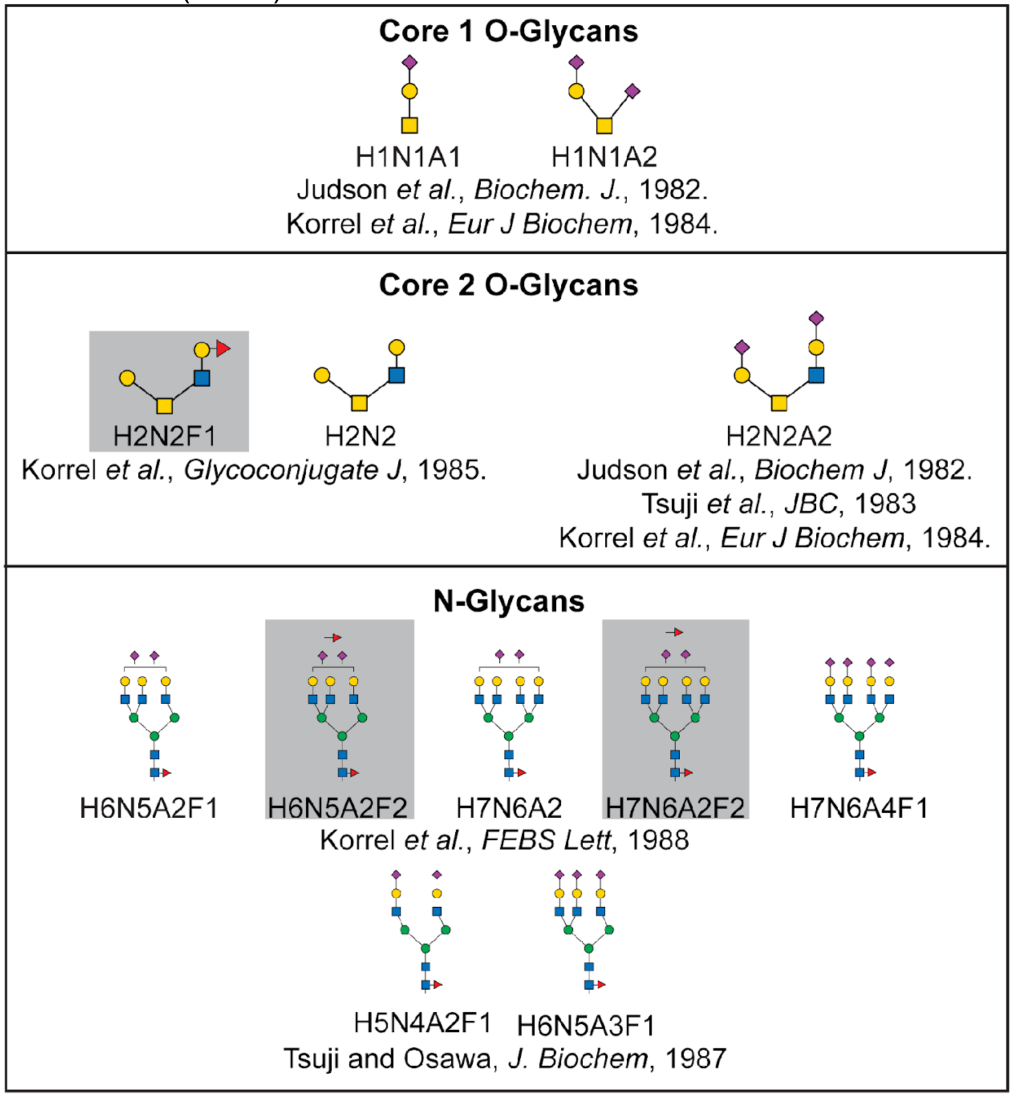

**Figure S3. Anion exchange chromatography purification of GPIbα ectodomain.** A) Full Ultraviolet (UV) absorbance at 280 nm *vs.* elution volume plot for the purification. B) Enlarged plot of the bracketed region in (A). GPIbα-containing fractions of highest purity eluted between 16-22 mL.

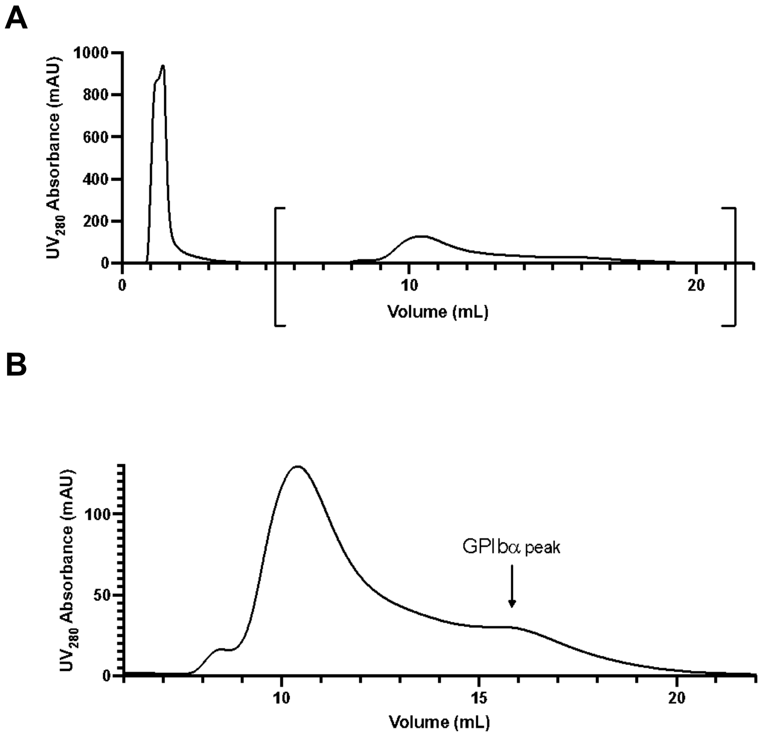

**Figure S4. SDS-PAGE and anti-GPIbα blot analysis of GPIbα purification.** Samples were analyzed by SDS-PAGE and Coomassie total protein stain as well as by anti-GPIbα western blot at each step in the purification protocol. The samples analyzed were 1) supernatant following platelet sonication and incubation at 37 °C (crude), 2) after WGA lectin column (post-WGA), and 3) after anion exchange chromatography (post-anion exchange). The final preparation of GPIbα appeared as a single band on Coomassie-stained gel.

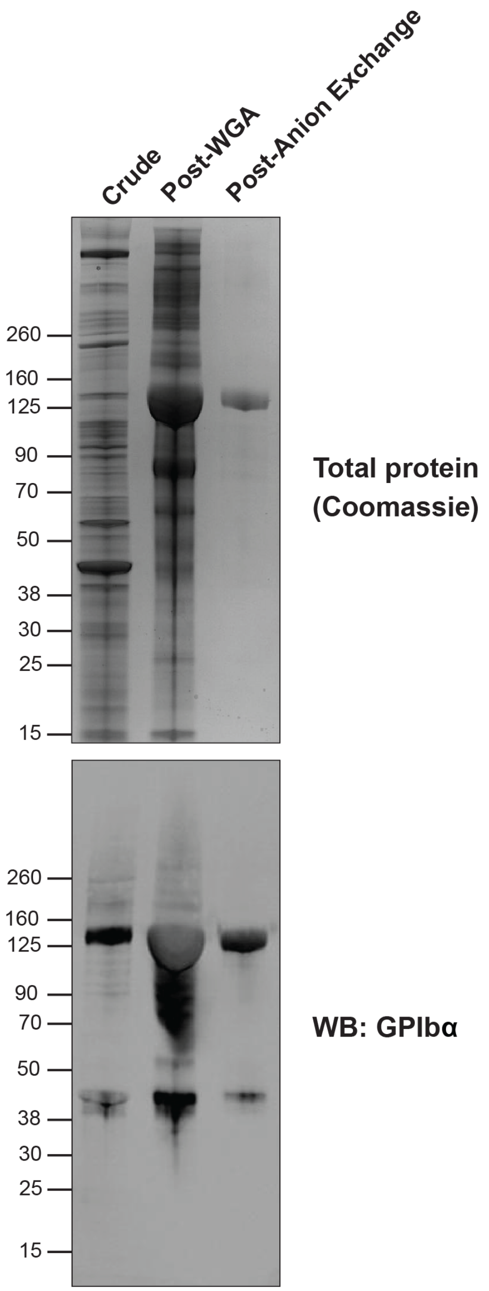

**Figure S5. Glycosyltransferase B converts GPIbα H antigens into B antigens.** GPIbα from type O platelets was treated with the recombinant human glycosyltransferase B and UDP-galactose. Immunoblotting of enzyme-treated samples showed strong anti-B antigen reactivity, while untreated samples were negative for any B antigen reactivity.

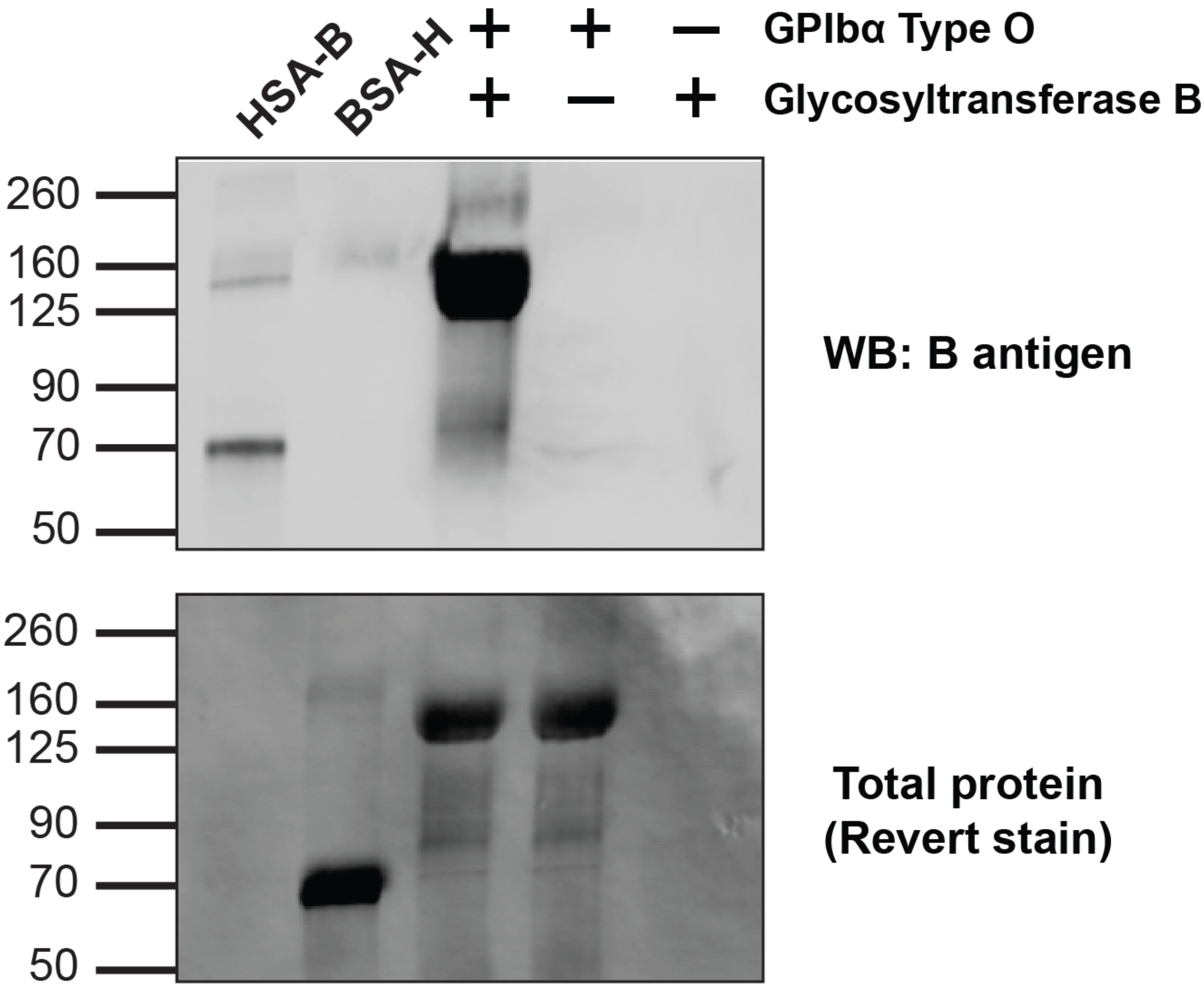

**Figure S6. PNGase F treatment of type A GPIbα does not eliminate A antigens.** GPIbα from type A platelets was treated with GalNAcEXO to remove the GalNAc that is the A antigen immunodominant epitope. This treatment completely abolished anti-A antigen reactivity. When GPIbα was treated with PNGase F to remove N-glycans, no significant change in anti-A antigen reactivity was observed, suggesting that the A antigen epitope is not restricted to N-glycans. Removal of N-glycans reduced SNA-biotin binding, suggesting that N-glycans contain sialic acid.

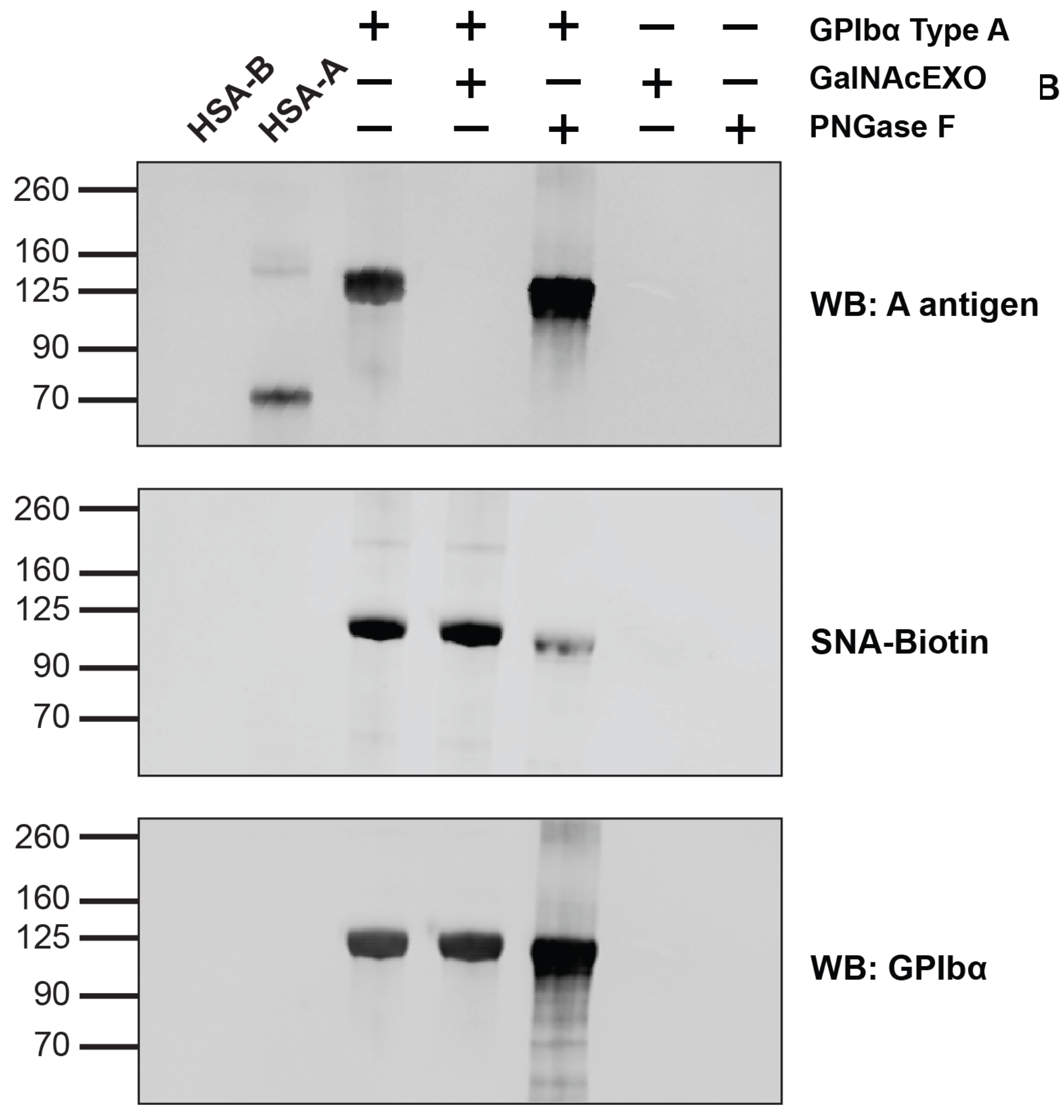

**Figure S7. PNGase F treatment of type B GPIbα does not eliminate B antigens.** GPIbα from type B platelets was treated with α-galactosidase to remove the Gal that is the B antigen determinant, and this treatment completely abolished anti-B antigen reactivity. When GPIbα was treated with PNGase F to remove N-glycans, no significant change in anti-B antigen reactivity was observed, indicating that the B antigen epitope is not restricted to N-glycans. Removal of N-glycans also reduced SNA-biotin binding, suggesting that N-glycans contain sialic acid.

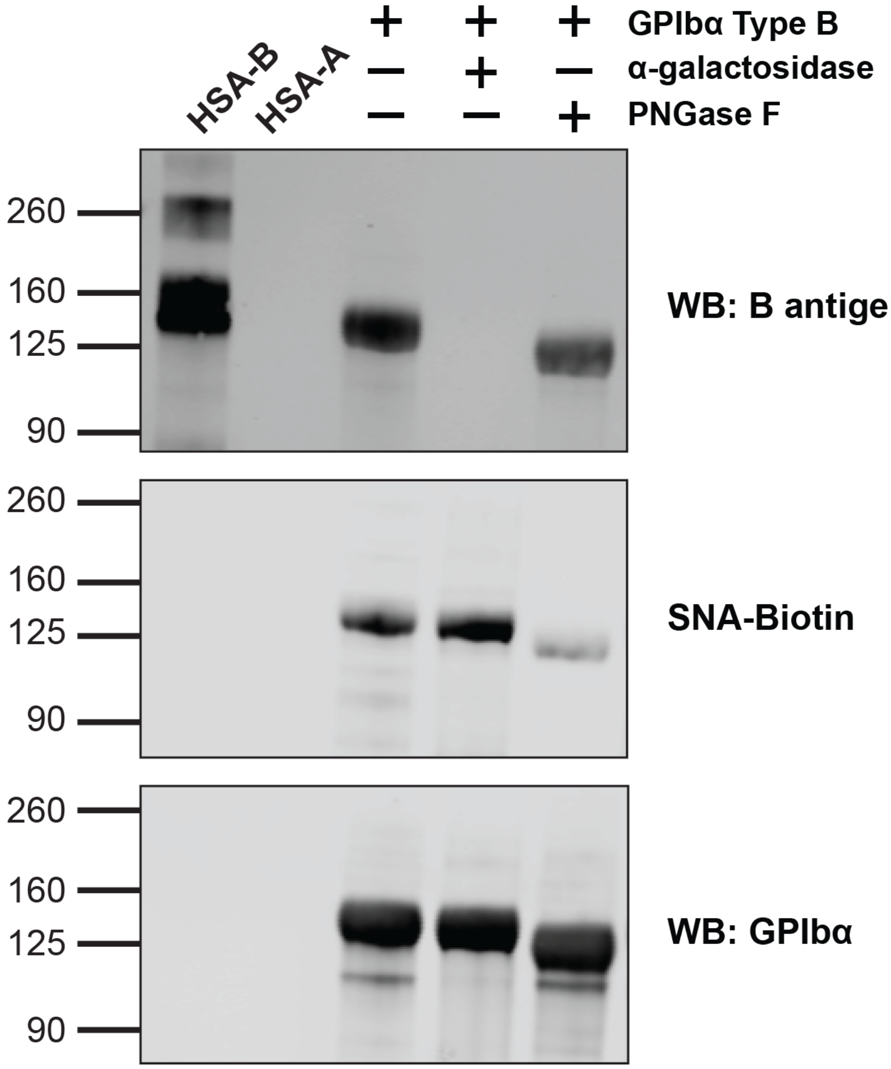

**Figure S8. PNGase F de-N-glycosylates fetuin.** Fetuin was incubated with PNGaseF**,** and the reaction was analyzed by SDS-PAGE. Following PNGaseF treatment, the molecular weight of fetuin is decreased due to N-glycan removal.

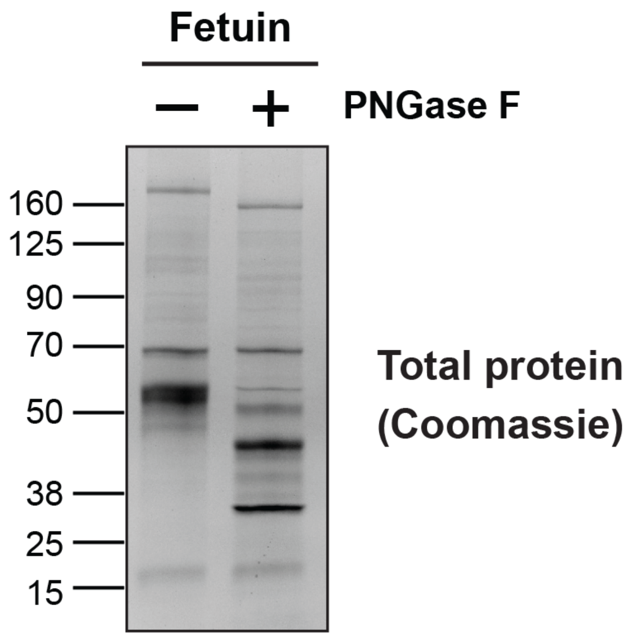

**Figure S9. Sialic acid quantification by DMB assay.** Signal intensity *vs.* retention time plot for reversed phase ultra-performance liquid chromatography, fluorescence (RP-UPLC-FL) to determine sialic acid content of GPIbα glycans from human blood type O platelets (n=3). Two common types of sialic acid were measured: *N*-acetyl-neuraminic acid (Neu5Ac) and *N*-glycolyl-neuraminic acid (Neu5Gc). Neu5Ac is the most common sialic acid in humans. Humans do not have the capacity to biosynthesize Neu5Gc. Sialic acid content was determined by comparison of peak integration values with known amounts of Neu5Ac and Neu5Gc standards. Neu5Gc was not detected in GPIbα samples.

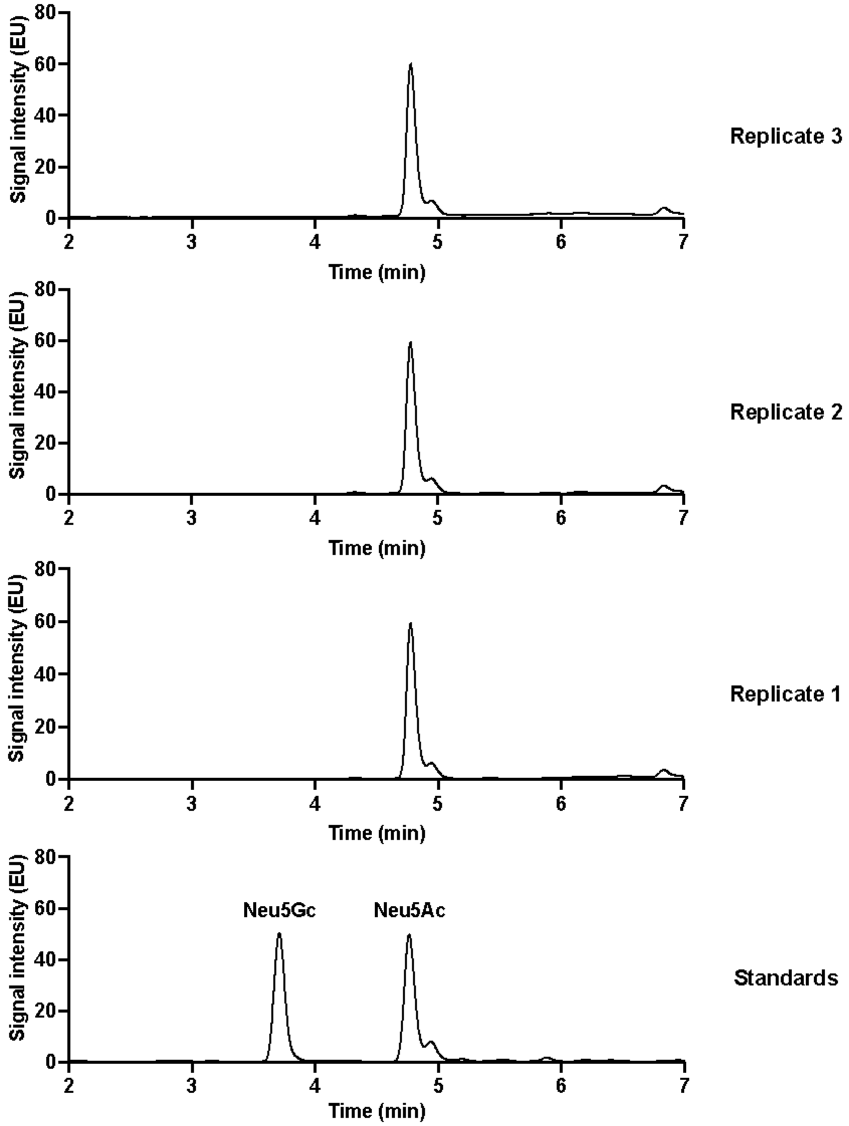

**Figure S10. Monosaccharide analysis by high-performance anion-exchange chromatography with pulsed amperometric detection (HPAEC-PAD).** Signal intensity *vs.* retention time plot for HPAEC-PAD to determine monosaccharide content of GPIbα glycans from blood type O platelets (n=3). Monosaccharide content was quantified by comparison of peak integration values with known amounts of standards of L-fucose, D-galactosamine, D-glucosamine, D-galactose, D-glucose, and D-mannose.

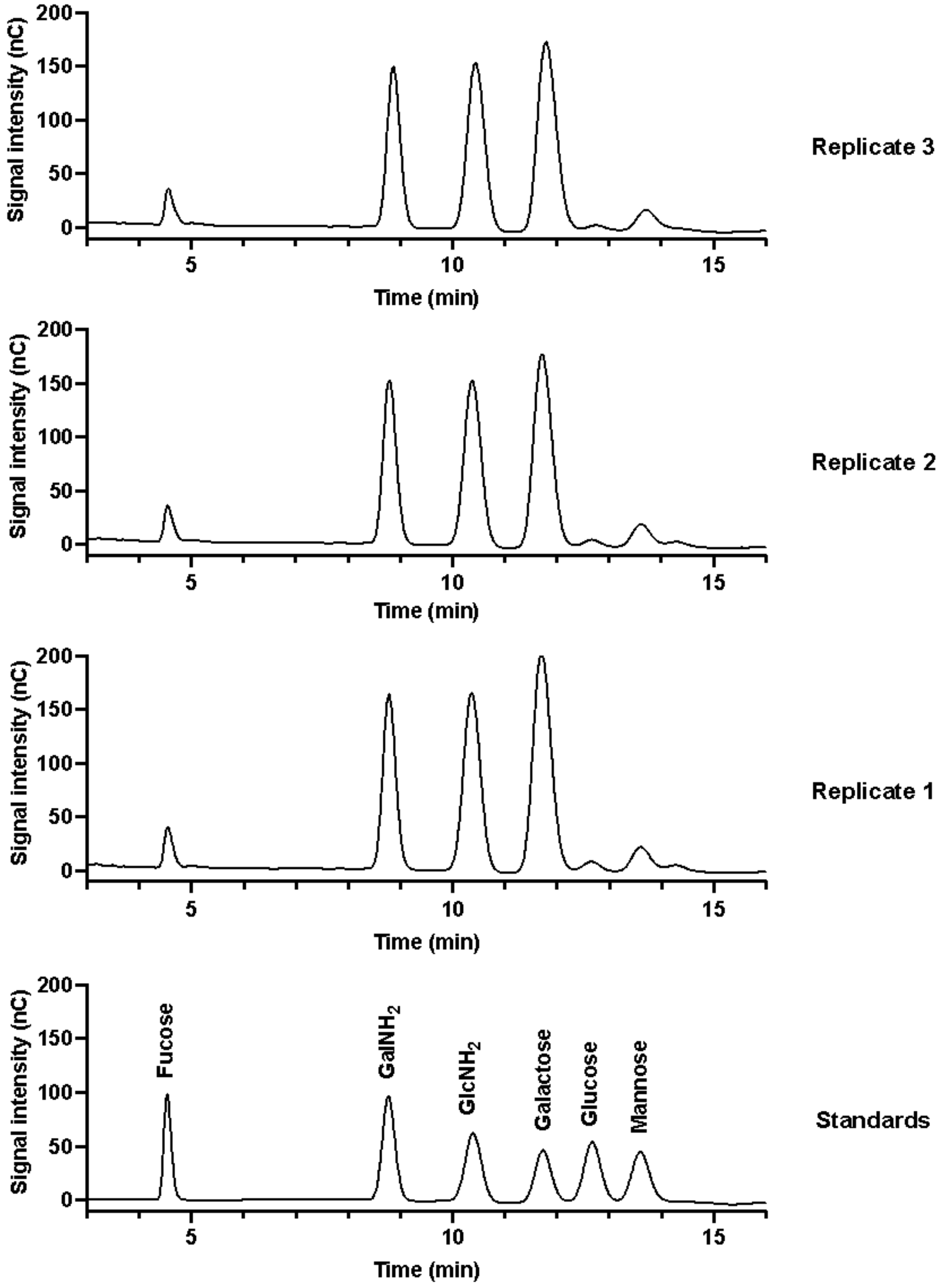

**Figure S11. O-glycans identified in glycomics analysis of GPIbα.** Glycomics analysis revealed 8 O-glycan structures that were > 3% abundance in the full MS profile. Each of these O-glycans was localized to one or more glycosites in the glycoproteomics analysis. H, hexose; N, *N*-acetylhexosamine; A, *N*-acetylneuraminic acid; F, fucose.

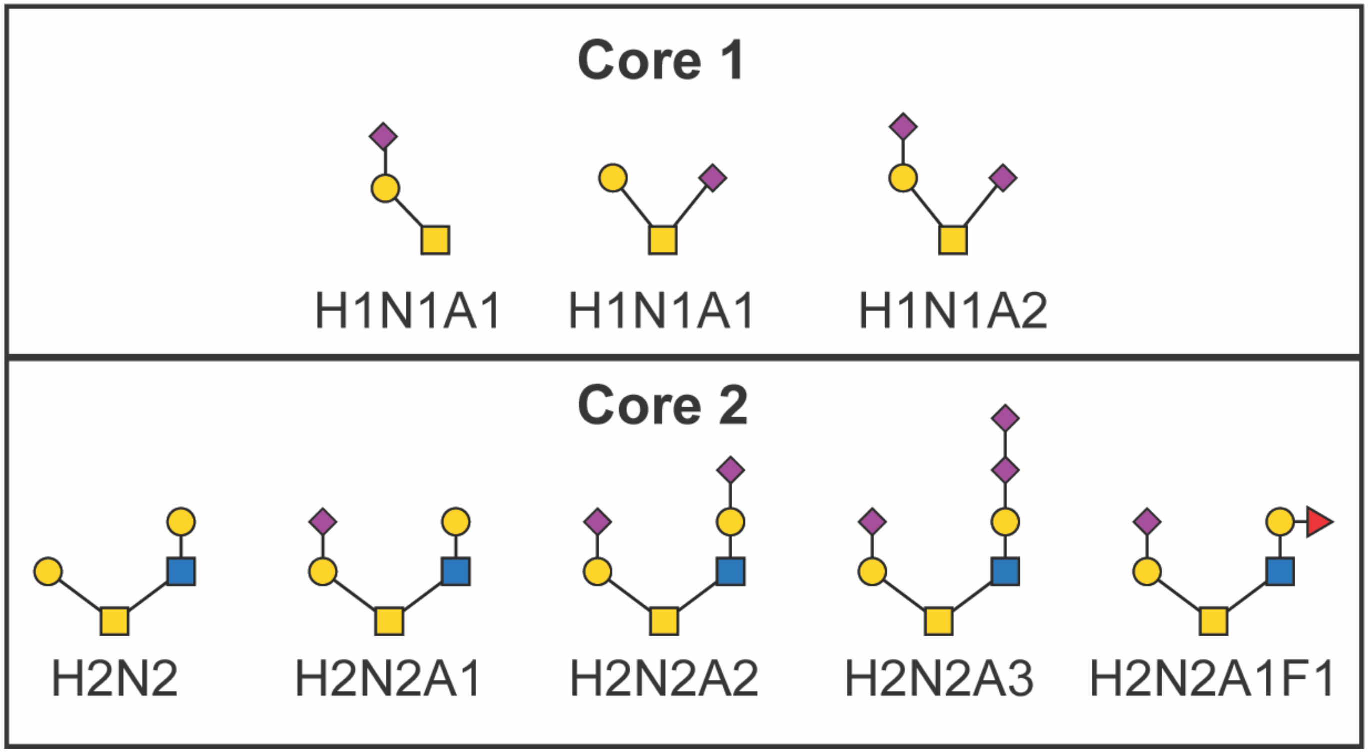

**Figure S12. MS$^3$ fragmentation of a Lewis antigen standard.** H and Lewis type antigens are trisaccharides comprised of Fuc, Gal, and GlcNAc. In the H antigen, Fuc is linked to Gal. In Lewis type antigens, Fuc is linked to GlcNAc. These structures can be discriminated by MS$^3$ fragmentation of the m/z = 660 trisaccharide. This figure shows the MS$^3$ fragmentation data for the Lewis antigen trisaccharide m/z 660, which generates diagnostic ions at *m/z* 456, 329, and 259. This contrasts with the MS$^3$ fragment ions from the H antigen m/z = 660 trisaccharide which are *m/z* 503, 433, and 415 (Figure 5B, Figure S16).

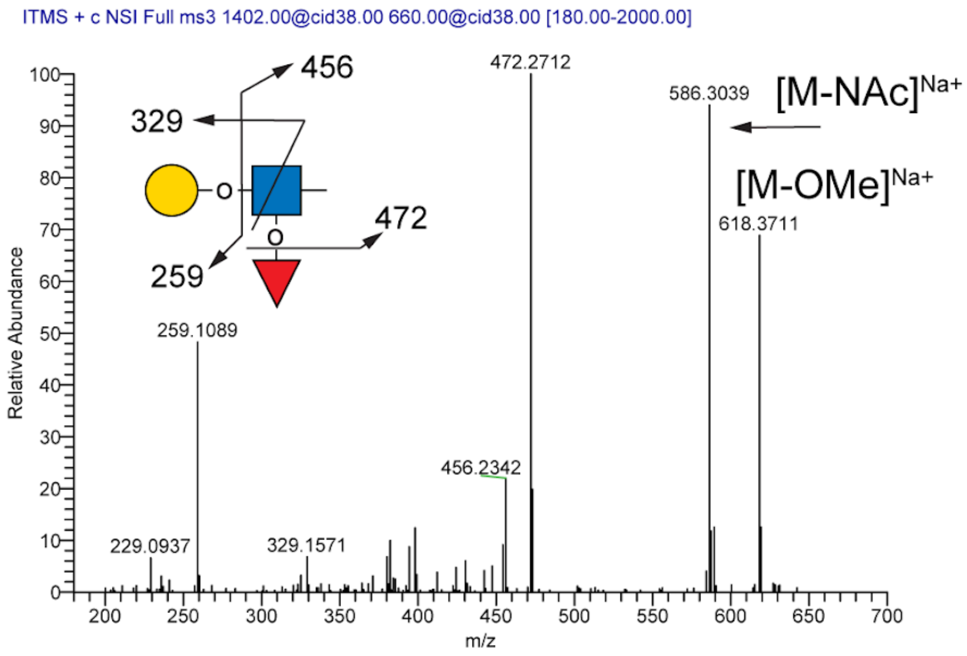

**Figure S13. O-glycans that carry ABH antigens identified in glycomics analysis of GPIbα.** The gray box indicates an O-glycan that was identified in previous analyses, as shown in Figure S2. The glycans with bolded monosaccharide composition labels are those that were site-localized to one or more glycosites in the glycoproteomics analysis. H, hexose; N, *N*-acetylhexosamine; A, *N*-acetylneuraminic acid; F, fucose.

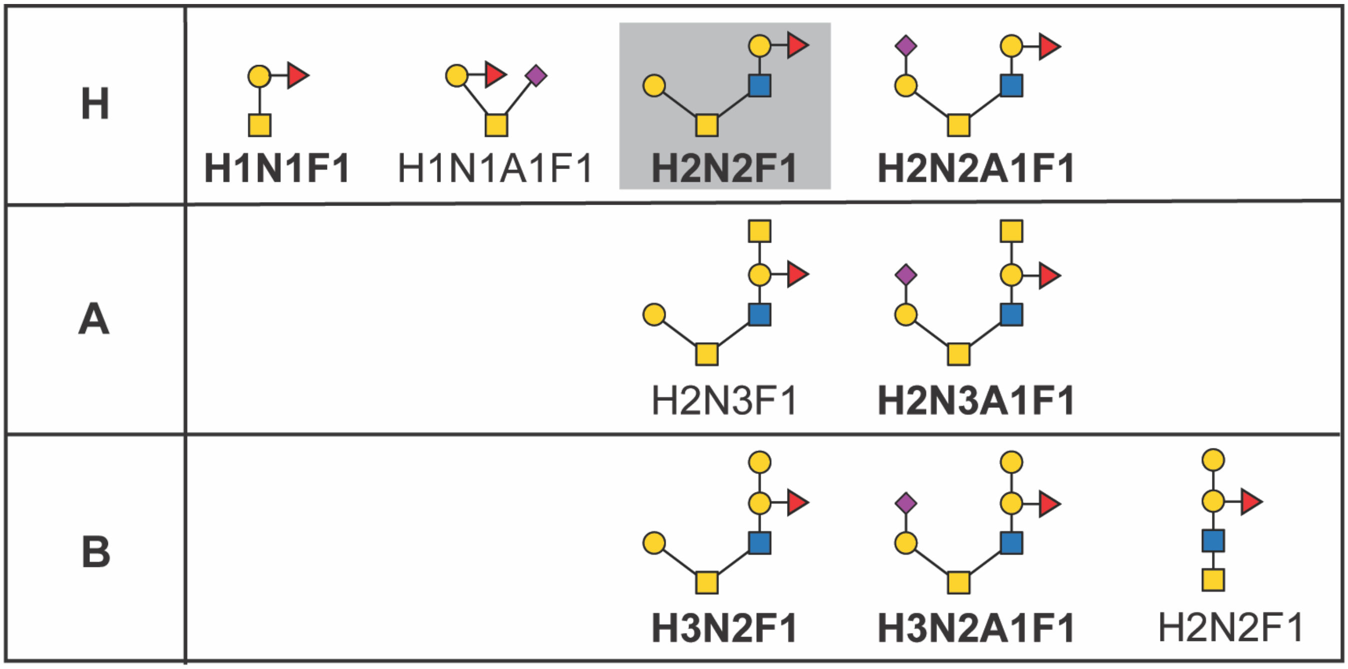

**Figure S14. Type B GPIbα carries B antigen on core 2 type O-glycan.** Type B GPIbα O-glycomics analysis revealed an O-glycan with m/z = 1361. $MS^2$ analysis generates a fragment ion with m/z = 864 consistent with the B antigen tetrasaccharide. $MS^3$ fragmentation of the m/z = 864 generated the expected diagnostic ions for B antigen (m/z = 707, 676, 637, and 619).

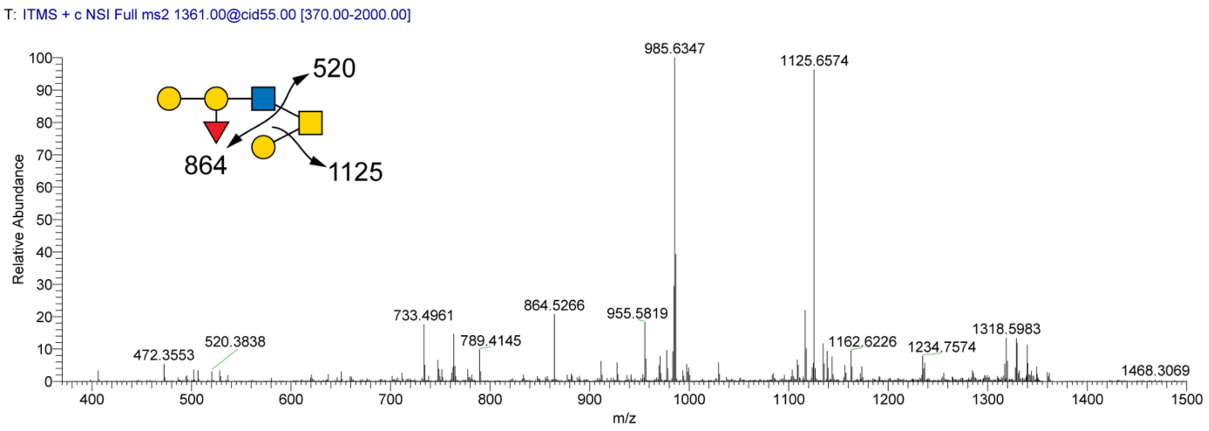


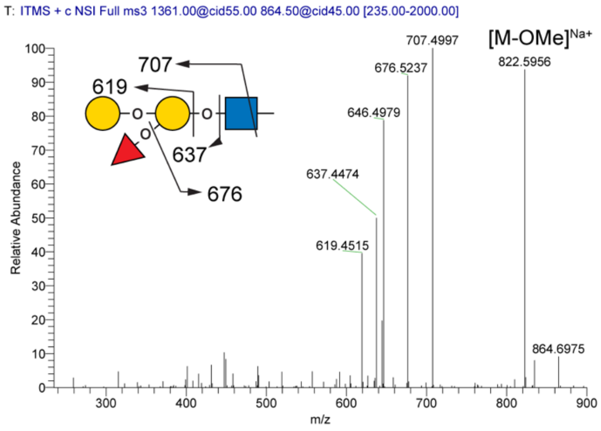


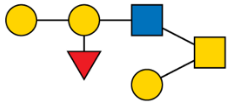

**Figure S15. N-glycans identified in glycomics analysis of GPIbα.** Glycomics analysis revealed 18 N-glycan structures that were > 3% abundance in the full MS profile.

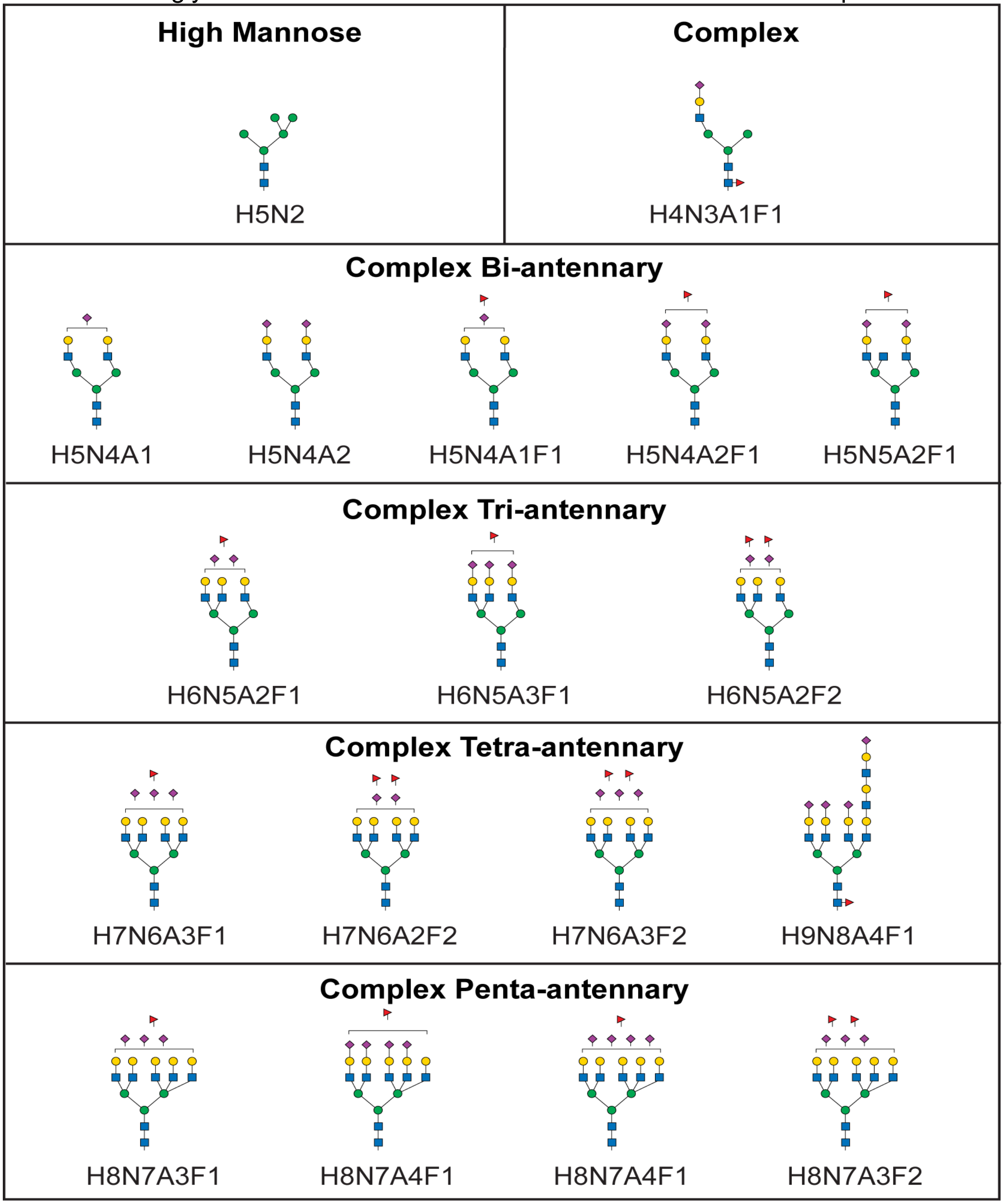

**S16. Type O GPIbα carries H antigen on a tetra-antennary complex N-glycan.** Type O GPIbα N-glycomics analysis revealed an N-glycan with m/z = 1482. MS$^2$ analysis generated a fragment ion with m/z = 660 consistent with the H antigen trisaccharide. On MS$^3$ fragmentation, this generated the expected diagnostic ions for H antigen (m/z = 503, 433, and 415), and not those expected for Lewis-type antigens.

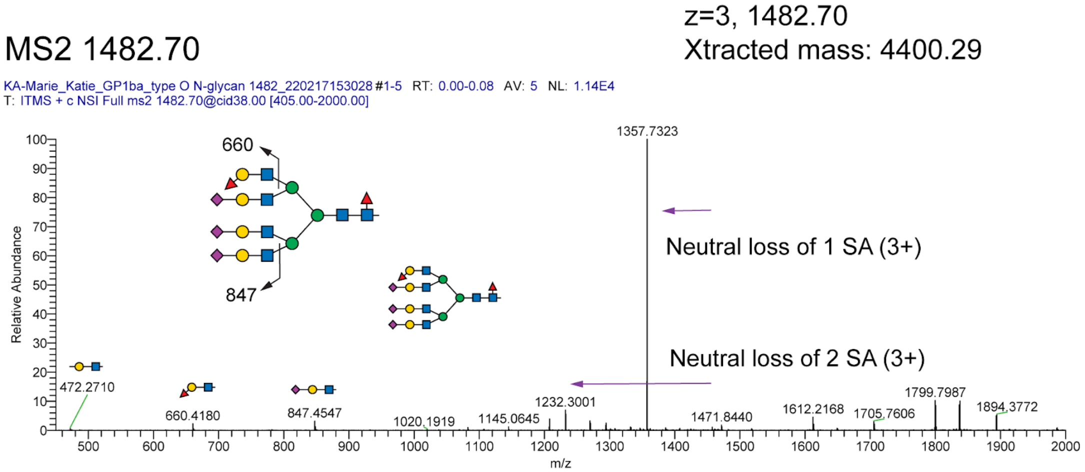


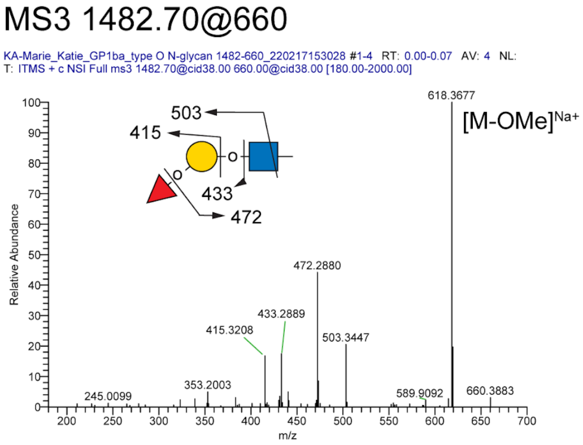

**Figure S17. N-glycans that carry ABH antigens identified in glycomics analysis of GPIbα.** The gray box indicates the N-glycans that were identified in previous analyses, as shown in Figure S2. H, hexose; N, *N*-acetylhexosamine; A, *N*-acetylneuraminic acid; F, fucose.

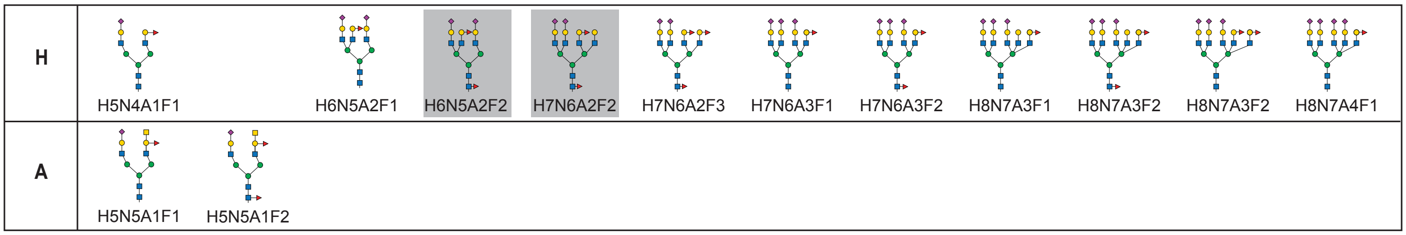

**Figure S18. Type A GPIbα carries both A and H antigen on bi-antennary complex N-glycans.** Type A GPIbα N-glycomics analysis revealed an N-glycan with m/z = 1524. $MS^n$ analysis suggests that this represents two isobaric N-glycans; one carrying the H antigen and one carrying the A antigen. $MS^2$ analysis generates: 1) a fragment ion with m/z = 660 consistent with the H antigen trisaccharide and 2) a fragment ion with m/z = 905 consistent with the A antigen tetrasaccharide. $MS^3$ fragmentation of the m/z = 990 generated the expected diagnostic ions for A antigen (m/z = 748, 717, 678, and 646).

MS2 1524.50

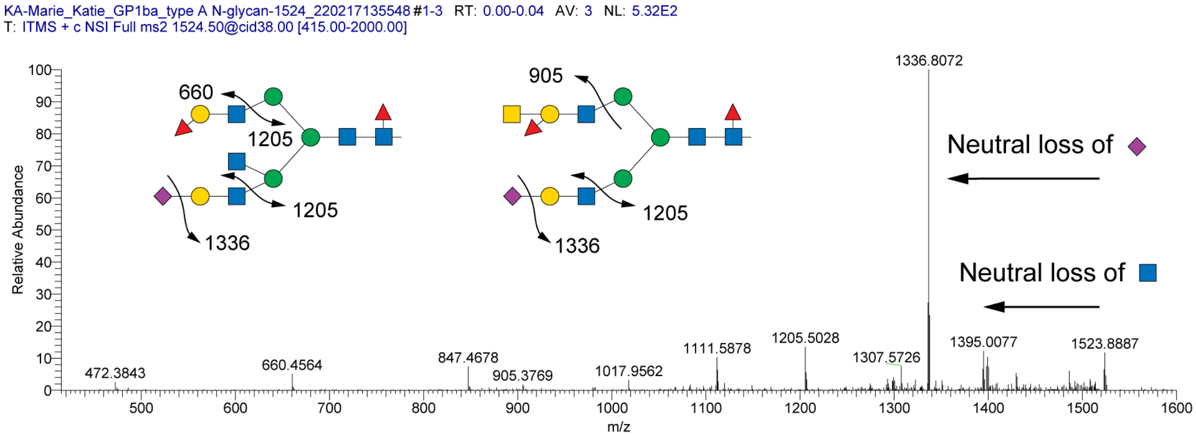


MS3 1524.50@905

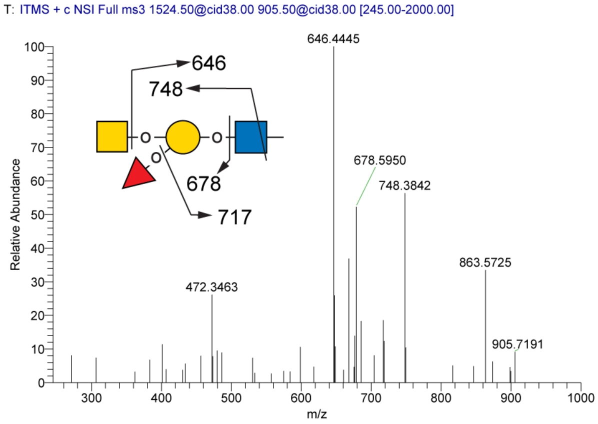

#### III. Supplemental Tables

**Table S1. Domain organization of the GPIbα ectodomain.** The mucin domain amino acids are defined based on our "mucin candidacy algorithm".(*22*)

| Amino Acids | Domain/Feature |
|---|---|
| 1-303 | Von Willebrand Factor ligand-binding domain |
| 304-458 | Macroglycopeptide domain |
| 371-460 | Mucin domain |
| 459-525 | Mechanosensory domain |
| 506-507 | ADAM17 cleavage site |

**Table S2. GPIbα ectodomain glycosites and glycan structures determined previously and in this analysis.** Mucin domain amino acids are colored red. When more than one amino acid is listed in a row, this indicates that, due to the tandem repeat sequences, the glycosite could not be assigned to a particular repeat. The "recombinant protein" refers to the human protein recombinantly expressed in a mouse myeloma cell line. The "platelet protein" refers to the protein purified from human paltelets. ND, no data (N-glycosite identification was not performed for the recombinant protein). H, hexose; N, *N*-acetylhexosamine; A, *N*-acetylneuraminic acid; F, fucose.

| Amino Acid | NetGlyc Predicted Glycosite | Previously Experimentally Determined Glycosite | Glycosite Identified Recombinant Protein | Glycosite Identified Platelet Protein | **Glycan Structures Identified Platelet Protein** | | | | |
|---|---|---|---|---|---|---|---|---|---|
| | | | | | Tn | T | ABH | Neu5Ac (Non-ABH) | Other |
| **O-glycosites** | | | | | | | | | |
| S23 | - | - | + | - | - | - | - | - | - |
| S27 | - | - | + | - | - | - | - | - | - |
| T39 | - | - | + | - | - | - | - | - | - |
| S55 | - | - | + | - | - | - | - | - | - |
| T154 | - | - | + | - | - | - | - | - | - |
| S170 | - | - | + | - | - | - | - | - | - |
| T192 | - | - | + | - | - | - | - | - | - |
| T256 | + | - | + | - | - | - | - | - | - |
| S257 | + | - | + | - | - | - | - | - | - |
| S261 | - | - | + | - | - | - | - | - | - |
| S267 | - | - | + | - | - | - | - | - | - |
| T282 | - | - | + | - | - | - | - | - | - |
| T289 | - | - | + | - | - | - | - | - | - |
| T308 | + | + | + | + | - | - | - | H1N1A2<br>H2N2A2 | - |
| T310 | + | - | + | - | - | - | - | - | - |
| T316 | + | - | + | + | - | - | - | H1N1A1 | - |
| T320 | + | - | + | + | - | - | - | H2N2A2 | - |
| T321 | - | - | + | + | - | - | H: H2N2A1F1 | H2N2A1<br>H2N2A2<br>H2N2A3 | H2N2 |
| S328 | + | - | + | + | - | - | - | H1N1A1 | - |
| S330 | + | - | + | - | - | - | - | - | - |
| T331 | - | - | + | + | - | - | - | H2N2A1 | - |
| S333 | - | - | + | - | - | - | - | - | - |
| S336 | - | - | + | - | - | - | - | - | - |
| S340 | + | - | + | + | - | + | - | H2N2A2 | H2N2 |
| S341 | + | - | + | + | - | + | - | H1N1A1<br>H1N1A2<br>H2N2A1<br>H2N2A2 | - |
| T345 | + | - | + | + | - | - | - | H1N1A1<br>H2N2A1<br>H2N2A2 | - |
| S348 | + | - | - | - | - | - | - | - | - |
| T349 | + | - | - | - | - | - | - | - | - |
| T353 | + | - | + | + | - | - | - | H1N1A1 | - |
| T354 | + | - | + | + | - | - | - | H1N1A2 | - |

| | | | | | | | | | |
|---|---|---|---|---|---|---|---|---|---|
| | | | | | | | | H2N2A2 | |
| T360 | + | - | + | + | - | - | - | H2N2A2 | - |
| T364 | + | - | + | + | - | - | H: H2N2A1F1<br>A: H2N3A1F1 | H2N2A1<br>H2N2A2<br>H1N1A2 | H2N2 |
| S369 | + | - | + | + | - | + | - | - | - |
| T371 | + | - | + | + | - | - | H: H2N2F1<br>H: H2N2A1F1 | H1N1A1<br>H1N1A2<br>H2N2A1 | H2N2 |
| S373 | + | - | + | + | - | + | - | H1N1A1<br>H1N1A2 | - |
| T375 | + | - | + | - | - | - | - | - | - |
| S378 | + | - | - | - | - | - | - | - | - |
| T379 | + | - | - | + | - | - | - | H1N1A1 | - |
| T380 | + | - | + | + | + | + | H: H2N2F1<br>A: H1N2F1 | H1N1A1 | H2N2 |
| T383 | + | - | + | + | + | + | - | - | H2N2 |
| S385 | + | - | + | + | + | - | - | - | - |
| T387 | + | - | + | + | + | + | - | - | - |
| T388 | + | - | + | + | + | - | - | - | N2 |
| S389 | + | - | + | + | + | - | H: H2N2A1F1 | H2N2A1 | - |
| T400 | + | - | - | + | + | + | - | H2N2A1 | - |
| T401 | + | - | + | + | - | + | - | - | - |
| T405 | + | - | + | + | - | + | - | - | - |
| S407;<br>420;<br>433 | + | - | - | + | + | + | H: H1N1F1 | H2N2A2 | - |
| T409;<br>422;<br>435;<br>448 | + | - | + | + | + | - | - | H1N1A1 | - |
| T410;<br>423;<br>436;<br>449 | + | - | - | + | + | + | - | - | - |
| T414;<br>427;<br>440 | + | - | - | + | + | - | - | - | - |
| S415;<br>428;<br>441 | + | - | + | + | + | + | - | H2N2A1<br>H2N2A2 | - |
| S420;<br>433;<br>446 | + | - | + | + | + | - | - | H1N1A1 | - |
| T453 | - | - | - | + | - | - | - | H1N1A1<br>H1N1A2<br>H2N2A1<br>H2N2A2<br>H2N2A3 | |
| T457 | + | - | + | + | - | - | - | H1N1A1 | - |
| T460 | + | - | + | + | - | + | - | - | - |
| S461 | + | - | + | + | - | + | - | H2N2A2 | - |
| T463 | + | - | + | + | - | - | H: H2N2F1<br>H: H2N2A1F1 | H1N1A1<br>H1N1A2<br>H2N2A1<br>H2N2A2 | H2N2 |
| S467 | + | - | + | + | - | - | - | H1N1A1<br>H2N2A1<br>H2N2A2 | H2N2 |
| T469 | + | + | + | + | - | - | H: H2N2A1F1<br>A: H2N3A1F1<br>B: H3N2A1F1 | H1N1A1<br>H1N1A2<br>H2N2A1<br>H2N2A2 | - |
| S470 | + | + | + | - | - | - | - | - | - |
| T473 | + | - | + | + | - | - | H: H2N2A1F1 | H1N1A1<br>H2N2A2<br>H2N2A1 | H2N2 |
| S476 | + | - | + | - | - | - | - | - | - |
| T477 | + | - | + | + | - | - | - | H1N1A1 | - |
| T480 | + | + | + | + | - | - | H: H1N1F1 | H1N1A1 | - |
| T481 | + | + | + | + | - | - | H: H2N2F1<br>H: H2N2A1F1 | H1N1A1<br>H2N2A1 | H1N1<br>H2N2 |
| T482 | + | + | + | + | - | + | H: H1N1F1<br>H: H2N2A1F1<br>H: H2N2F1<br>A: H2N3A1F1 | H1N1A1<br>H2N2A1<br>H2N2A2 | H2N2 |
| S486 | + | - | + | + | - | - | H: H2N2F1 | H1N1A1<br>H1N1A2 | - |

| | | | | | | | | | |
|---|---|---|---|---|---|---|---|---|---|
| S490 | + | - | + | + | - | + | - | H1N1A1 | - |
| T491 | + | + | + | + | - | + | H: H2N2F1<br>H: H2N2A1F1<br>H: H1N1F1<br>A: H2N3A1F1<br>B: H3N2A1F1 | H1N1A1<br>H1N1A2<br>H1N2A1<br>H2N2A1<br>H2N2A2<br>H2N2A3 | H2N2 |
| T494 | + | + | + | + | + | + | H: H1N1F1<br>H: H2N2F1<br>H: H2N2A1F1<br>B: H3N2F1 | H1N1A1<br>H1N1A2<br>H2N2A1<br>H2N2A2 | H2N2 |
| **N-Glycosites** | | | | | **N-glycans** | | | | |
| N37 | + | + | ND | + | H5N4A2F1<br>H6N5A2F1<br>H6N5A3F1<br>H6N5A2F2<br>H6N5A3F1<br>H6N6A1F2<br>H6N6A1F7<br>H6N6A1F3<br>H6N6A3F1<br>H7N6A1F4<br>H7N6A2F2<br>H7N6A3F1<br>H7N6A3F2<br>H7N6A4F1<br>H8N7A3F2<br>H8N7A4F1 | | | | |
| N175 | + | + | ND | - | - | | | | |
| N362 | + | - | ND | - | - | | | | |
| N398 | + | - | ND | - | - | | | | |

**Table S3. Platelet donor ABO blood group determination.** Platelet donor ABO blood group determination was made based on results from the forward and reverse type as indicated by the table below. A “+” indicates agglutination and “-“ indicates no agglutination. RBCs, red blood cells.

| | Forward Type | | | Reverse Type | | |
|---|---|---|---|---|---|---|
| **Reagent**<br><br>**Donor** | Anti-A antibody RBCs | Anti-B antibody RBCs | Anti-A,B antibody RBCs | A1 RBCs<br><br>Serum/Plasma | B RBCs<br><br>Serum/plasma | **Assigned Blood Group** |
| Agglutination result | + | - | + | - | + | A |
| | - | + | + | + | - | B |
| | - | - | - | + | + | O |

**Table S4. Representative GPIbα ectodomain purification yields.** Five units of apheresis platelets with a total combined volume of 1219 mL were subjected to sonication and incubation at 37 °C, WGA lectin chromatography, and anion exchange chromatography to purify the soluble ectodomain of GPIbα. Following each purification step, total protein concentration was measured by UV absorbance at 280 nm.

| Purification Step | Total protein yield |
|---|---|
| Sonication supernatant | 1260 mg |
| WGA lectin column | 4.9 mg |
| Anion exchange chromatography | 350 μg |
| Starting material: 1219 mL apheresis platelets | |
| Yield: 0.3 μg / mL of platelets (70 μg / unit) | |

**Table S5. Gradient used for UPLC separation of DMB-sialic acid.**

| Time (min) | Solvent A | Solvent B |
|---|---|---|
| 0 | 100 | 0 |
| 10 | 92 | 8 |
| 11 | 100 | 0 |
| 18 | 100 | 0 |

**Table S6. Gradient used for HPAEC-PAD separation of monosaccharides.**

| Time (min) | Solvent A | Solvent B | Solvent C |
|---|---|---|---|
| 0 | 81 | 19 | 0 |
| 23 | 0 | 15 | 85 |
| 35 | 81 | 19 | 0 |
| 50 | 81 | 19 | 0 |

**Table S7. Monosaccharide composition of GPIbα ectodomain**. GPIbα isolated from human platelets from blood type O donors (n=3) was determined by HPAEC-PAD (for Gal, GlcNac, GalNAc, mannose, fucose, and glucose) and DMB-assay followed by RP-UPLC-FL (for sialic acid). Molar amounts of each monosaccharide were divided by the molar quantity of GPIbα ectodomain that was analyzed to calculate molar ratios.

| Blood Type O | Neu5Ac:GPIbα (molar ratio) | Galactose:GPIbα (molar ratio) | GlcNAc:GPIbα (molar ratio) | GalNAc:GPIbα (molar ratio) | Mannose:GPIbα (molar ratio) | Fucose:GPIbα (molar ratio) | Glucose:GPIbα (molar ratio) |
|---|---|---|---|---|---|---|---|
| **Replicate 1** | 58.5 | 54.4 | 32.6 | 21.0 | 6.7 | 5.0 | 2.0 |
| **Replicate 2** | 58.6 | 54.9 | 33.2 | 20.6 | 6.6 | 5.0 | 1.7 |
| **Replicate 3** | 58.7 | 60.2 | 33.9 | 21.8 | 6.9 | 5.3 | 2.2 |
| **Mean ± std dev** | 58.6 ± 0.1 | 56.5 ± 3.2 | 33.2 ± 0.7 | 21.1 ± 0.6 | 6.7 ± 0.1 | 5.1 ± 0.2 | 2.0 ± 0.3 |

**Table S8. Glycan structures identified in glycoproteomics analysis of recombinant GPIbα ectodomain.** H = hexose, N = *N*-acetylhexosamine, A = *N*-acetylneuraminic acid, G = *N*-glycolylneuraminic acid, F= fucose
*Glycan identified only in sialidase treated sample.

| | Glycan # of Monosaccharides | | | | | |
|---|---|---|---|---|---|---|
| Amino Acid | 1 | 2 | 3 | 4 | 5 | 6 |
| S23 | - | - | H2N1* | - | H1N2A1F1 | - |
| S27 | N1* | - | - | - | - | - |
| T39 | - | - | - | - | H2N2F1 | H2N2A2 |
| S55 | - | - | - | - | H1N2F2 | - |
| T154 | - | - | - | - | H1N2A1F1 | - |
| S170 | - | - | - | - | H1N2A1F1 | - |
| T192 | - | - | - | H1N2F1 | - | - |
| T256 | N1 | N2 | - | - | H1N2A1F1* | - |
| S257 | - | N2 | - | - | H1N2A1F1* | - |
| S261 | N1* | N2 | - | H2N1F1<br>H2N2* | H1N2A1F1 | - |
| S267 | N1* | - | H1N2 | H1N1A1F1<br>H1N2F1*<br>H2N1F1 | - | - |
| T282 | - | - | - | - | - | H2N2A1F1* |
| T289 | - | - | H1N2* | - | - | - |
| T308 | N1* | H1N1 | H1N1A1<br>H1N1G1 | H1N1A2<br>H1N1G2<br>H1N2G1 | - | - |
| T310 | N1* | N1A1*<br>H1N1*<br>N2* | H1N1A1* | H1N1A2 | - | - |
| T316 | N1* | H1N1<br>N2* | H1N1A1* | H2N2* | - | - |
| T320 | N1* | H1N1* | H1N1A1<br>H1N1G1 | H1N1A2<br>H1N1G2<br>H2N2* | - | - |
| T321 | N1* | H1N1* | - | H1N1G2<br>H2N2* | - | - |
| S328 | N1 | H1N1* | - | H1N1G2<br>H1N2F1*<br>H2N2* | - | - |
| S330 | N1* | H1N1* | H1N2* | - | - | - |
| T331 | N1* | H1N1*<br>N1A1 | H1N2* | H1N1A2 | H1N2F2* | - |
| S333 | - | H1N1*<br>N1A1* | H2N1* | H2N2* | H1N2F2* | - |
| S336 | N1* | H1N1*<br>N1A1<br>N2* | - | - | - | - |
| S340 | N1* | H1N1*<br>N1A1<br>N2* | - | - | - | - |
| S341 | N1* | H1N1*<br>N2* | - | H2N2* | H2N2A1 | - |
| T345 | N1* | H1N1* | H1N2* | - | H1N2A1F1*<br>H1N2F2* | |
| T353 | - | H1N1* | - | H2N2* | - | - |
| T354 | N1* | H1N1* | - | - | - | - |
| T360 | N1* | H1N1* | - | - | - | - |
| T364 | N1* | N2* | - | H2N2* | - | - |

| | | | | | | |
|---|---|---|---|---|---|---|
| | | H1N1* | | H1N1A2 | | |
| S369 | N1* | H1N1*<br>N1A1* | H1N1F1* | H1N1A2 | - | - |
| T371 | N1* | H1N1*<br>N1A1* | H1N1F1<br>H1N1G1<br>H1N1A1 | H1N1A2 | H1N2F2* | - |
| S373 | N1* | H1N1<br>N1A1* | H1N1G1<br>H1N1A1<br>H1N1F1* | - | - | - |
| T375 | N1* | H1N1 | - | - | - | - |
| T380 | N1* | H1N1* | H2N1* | - | - | - |
| T383 | - | - | H2N1* | - | - | - |
| S385 | - | N1A1* | - | - | - | H2N2F2* |
| T387 | - | H1N1*<br>N1A1* | - | H2N2* | - | H2N2F2* |
| T388 | N1* | H1N1* | - | - | - | - |
| S389 | - | - | H1N1A1*<br>H1N1F1* | H1N1A2*<br>H1N2F1* | - | - |
| T401 | N1* | H1N1* | - | - | - | - |
| T405 | N1* | H1N1* | H1N2* | - | - | - |
| T409 | N1* | - | - | - | - | - |
| S415 | - | - | - | H2N2* | - | - |
| S420 | N1* | - | H2N1 | - | - | - |
| T457 | N1* | H1N1* | H2N1* | H2N2* | H1N2A1* | - |
| T460 | N1* | H1N1* | - | H2N2<br>H2N1F1* | - | - |
| S461 | - | H1N1* | - | - | - | - |
| T463 | N1* | N2*<br>H1N1* | H1N2<br>H1N1A1* | - | H2N2F1* | - |
| S467 | N1* | H1N1* | H2N1* | - | - | H2N2F2* |
| T469 | N1* | H1N1* | H1N1G1 | H1N1G2<br>H1N1A2 | - | - |
| S470 | N1* | N2*<br>H1N1*<br>N1A1* | - | H1N2A1* | - | - |
| T473 | N1* | H1N1* | H1N1A1<br>H1N1G1 | H1N1A2<br>H1N1G2<br>H1N2F1<br>H2N1G1*<br>H2N2* | H2N2A1* | - |
| S476 | H1N1* | - | H2N1* | H1N2A1*<br>H1N2F1* | - | - |
| T477 | N1* | H1N1* | - | - | - | - |
| T480 | N1* | N2*<br>H1N1* | H1N1G1 | H1N1A2 | - | - |
| T481 | N1 | H1N1 | H2N1* | H1N1A1<br>H1N1G1<br>H1N1A2<br>H1N1A1F1<br>H2N1F1*<br>H2N2* | H1N2F2* | - |
| T482 | N1* | H1N1<br>N1A1 | H1N2<br>H1N1A1<br>H1N1F1<br>H1N1G1<br>H2N1* | H1N1A2<br>H1N1A1F1<br>H1N1G2<br>H1N1A2<br>H1N2F1<br>H2N2*<br>H2N1F1 | H2N1A2 | H1N2A3 |
| S486 | N1* | H1N1*<br>N1A1 | H2N1 | H1N1A1F1<br>H1N1A2 | H2N1A2<br>H2N1A1F1 | - |

| | | | | | | |
|---|---|---|---|---|---|---|
| | | | | H1N2F1<br>H2N2 | H1N2F2* | |
| S490 | N1 | N2*<br>H1N1<br>N1A1 | H1N1A1<br>H1N1F1<br>H1N1G1<br>H2N1 | H1N1G2<br>H2N1F1<br>H2N1G1 | H2N1A2<br>H2N2F2 | - |
| T491 | N1* | H1N1<br>N1A1 | H1N1A1<br>H1N1F1<br>H1N1G1<br>H1N2* | H1N1A2<br>H1N1A1F1<br>H1N1G2<br>H2N1F1 | H2N1A1F1<br>H2N1A2 | H2N2F2 |
| T494 | N1 | H1N1<br>N1A1 | H1N1A1<br>H1N1F1<br>H1N1G1<br>H2N1 | H1N1A1F1<br>H1N1A2<br>H1N1G2<br>H1N2A1<br>H1N2G1<br>H2N1F1<br>H2N1G1*<br>H2N2 | H2N1A1F1<br>H1N1A3<br>H2N1A2 | H1N2A3<br>H2N2F2 |

#### IV. Supplemental References.
